# A Proxitome–RNA–capture Approach Reveals that Processing Bodies Repress Co–Regulated Hubs

**DOI:** 10.1101/2023.07.26.550742

**Authors:** Chen Liu, Andriani Mentzelopoulou, Ioannis H. Hatzianestis, Epameinondas Tzagkarakis, Vassilis Scaltsoyiannes, Xuemin Ma, Vassiliki A. Michalopoulou, Francisco J. Romero–Campero, Ana B. Romero–Losada, Panagiotis F. Sarris, Peter Marhavy, Bettina Bölter, Alexandros Kanterakis, Emilio Gutierrez–Beltran, Panagiotis N. Moschou

## Abstract

Cellular condensates are usually ribonucleoprotein assemblies with liquid– or solid–like properties. Because they lack a delineating membrane, the compositional determination of condensates is laborious. Here we set up a pipeline for proximity–biotinylation–dependent capture of RNA to investigate the RNA composition of the condensate in Arabidopsis known as the processing bodies (PBs). Using this pipeline together with *in situ* protein–protein interaction and RNA detection, *in silico*, and high–resolution imaging approaches, we studied PBs under normal and heat stress conditions. The composition of PBs in RNAs is much more dynamic than that of the total transcriptome. RNAs involved in cell wall development and regeneration, hormonal signaling, secondary metabolism/defense, and RNA metabolism were enriched in PBs. RNA binding proteins and liquid–to–solid phase transitions modulated specificity of RNA recruitment in PBs. Surprisingly, RNAs were sometimes recruited together with their encoded proteins. In PBs RNAs follow distinct fates, with small liquid-like PBs modulating RNA decay while larger ones storage. The size and properties of PBs are regulated by the actin polymerization cAMP receptor (SCAR)–WASP family verprolin homologous (WAVE) complex. SCAR/WAVE modulates signaling by shuttling RNAs between PBs and the translational machinery adjusting the ethylene signaling pathway. Heat stress leads to the storage of immunity–related RNAs in PBs by reducing PBs dynamics, suggesting why processes such as immunity malfunction under heat stress. In summary, we provide a method to identify RNAs in condensates which allowed us to reveal a mechanism for RNA fate regulation.

## Introduction

Intra– or intermolecular interactions between proteins and nucleic acids can promote their “demixing” from the bulk cytoplasmic pool to droplet–like ribonucleoprotein (RNP) condensates. RNP condensates can form through “liquid–liquid phase separation (LLPS)” usually driven by multivalent (i.e., many–low affinity binding sites) interactions between unstructured protein segments and/or nucleic acids (Beutel et al., 2019). Furthermore, LLPS is sensitive to concentration; only above a threshold (saturation concentration) do competent RNAs and proteins form or enter condensates (Alberti et al., 2019; Hatzianestis et al., 2023). Because many condensates form through LLPS they show properties of liquids but given enough time they can “solidify” whereby they show a high internal viscosity and reduced exchange of molecules (Jawerth et al., 2020). In non–plants, these “liquid–to–solid” transitions are functional, and regulate processes such as cell division, signaling, and immunity (Lasker et al., 2022).

The unstructured protein segments driving LLPS are low–complexity regions (LCRs), such as intrinsically disordered regions (IDRs) and prion–like domains (PrLDs) (Wiedner and Giudice, 2021). IDRs and PrLDs can have charged amino acids (such as arginine, lysine, and histidine) often punctuated by aromatic ones (Zaccara and Jaffrey, 2020). In addition, folded/structured segments of proteins can contribute to LLPS (Dignon et al., 2020). We know comparably little of what drives LLPS of RNA molecules.

Examples of RNP condensates are the evolutionarily conserved processing bodies (PBs), and stress granules (SGs) (Solis-Miranda et al., 2023). PBs and SGs numbers and sizes depend on conditions; furthermore, the size of SGs depends on small metabolic molecules (Chodasiewicz et al., 2022). Under heat stress (HS) PBs size can increase (Gutierrez-Beltran et al., 2015). In yeast, plants, and animals, PBs contain the RNA decapping, exosome, and deadenylase complexes, and RNA binding proteins (Liu et al., 2023). PBs have been associated with RNA decay, as they are enriched in the RNA decapping complex, comprising among others the plant Decapping 1 (DCP1), DCP2/Trident, as well as the DCP5, the scaffolding VARICOSE, and the exoribonuclease 4 (XRN4). XRN4 is a processive 5’–3’ exonuclease executing RNA decay by trimming RNAs decapped by the active pair DCP1–DCP2. Mutants of *DCP1*, *DCP2*, and *VARICOSE* are dwarf with trichome abnormalities and disorganized veins (Xu et al., 2006; Xu and Chua, 2009). Yet, XRN4 mutants do not show similar phenotypes, suggesting that PBs may execute additional functions or use additional ribonucleases. Although we know little about plant PBs, animal PBs not only degrade RNAs but can store them for conditional translation (Cairo et al., 2022). Yet, the mechanisms releasing RNAs from PBs for translation, as well as those modulating RNA fate in them are under intense scrutiny [e.g., (Blake et al., 2023)].

Studies on condensates’ functions would benefit from their isolation. As condensates lack a delineating membrane, their isolation is challenging due to their liquid nature (Liu et al., 2023). Nevertheless, in non–plants the catalog of proteins and RNAs associated with PBs has greatly increased in the past few years. Protein cataloging for PBs was recently done by proximity–dependent ligation of biotin (hereafter “PDL”) in non–plants (Youn et al., 2018) and plants (Liu et al., 2023). PDL harnesses covalent biotinylation of proteins interacting with or near–neighbors of a certain prey protein *in vivo*. Through PDL, we showed that the PBs component DCP1 co–condenses with the actin nucleating suppressor of the cAMP receptor (SCAR)–WASP family verprolin homologous (WAVE) complex in differentiating cells (Liu et al., 2023). This co-condensation takes place specifically at cell edges or vertices (the angular point where three cell edges meet) mainly at the transition and differentiation root zones. This “SCAR/WAVE–DCP1–axis” is important for proper development, but how it works remains unclear.

Inspired by the PDL application for the identification of the PBs proteome, we repurposed, here, PDL to isolate PBs–resident RNAs. We further show that large PBs are solidified and keep mainly translationally inert RNAs with their cognate proteins; interestingly, these RNAs function in similar pathways related to development and secondary metabolism, immunity, cell wall-wounding/regeneration, and hormonal signaling. On the contrary, smaller PBs show high liquidity and execute RNA decay. The SCAR/WAVE complex regulates the balance between small and large PBs, and by dissolving PBs, allows the release of stored RNAs or the reduction of their decay. We further show that the SCAR/WAVE–DCP1–axis modulates ethylene response, by regulating RNA fate.

## Results and Discussion

### A TurboID–RIP approach captures RNAs associated with PBs

The PDL with the biotin ligase BirA* has efficiently been used for RNPs purifications in animals (Leidal et al., 2020). Inspired by such applications, we assumed that the biotin ligase TurboID, which we and others have successfully used in plants (Arora et al., 2020), can also biotinylate proteins bound to RNAs. In turn, these biotinylated proteins could theoretically be cross–linked with their bound RNAs through formaldehyde. For this approach, referred to hereafter as “**<u>T</u>**urbo–**<u>R</u>**NA **I**mmuno**<u>p</u>**recipitation” (T–RIP; experimental approach detailed in **Fig. 1A** and Materials), we used Arabidopsis seedlings carrying transgenes encoding either a chimeric DCP1–TurboID–6x**<u>H</u>**is– 3x**<u>F</u>**LAG (*DCP1*–*TurboID–HF*) or the super folder–GFP under the 35S promoter [*GFP– TurboID–HF*, (Liu et al., 2023)]. *DCP1–TurboID–HF* could rescue the weak *dcp1–3* allele mutant phenotype [see below **Fig. 10** and (Martinez de Alba et al., 2015; Liu et al., 2023)], and showed sufficient biotinylation in immunoblots under the conditions used here (**Supplemental Fig. 1A**). However, we did not see sufficient biotinylation under the same conditions and various settings using the engineered and codon–optimized biotin ligase evolved from the soybean ascorbate peroxidase (“APEX2”, **Supplemental Fig. 1B –E**). This result contrasts with what has been observed in non–plant models where APEX2 biotinylates directly RNA and proteins (Fazal et al., 2019). We thus proceeded with the T–RIP approach.

**Figure 1.**
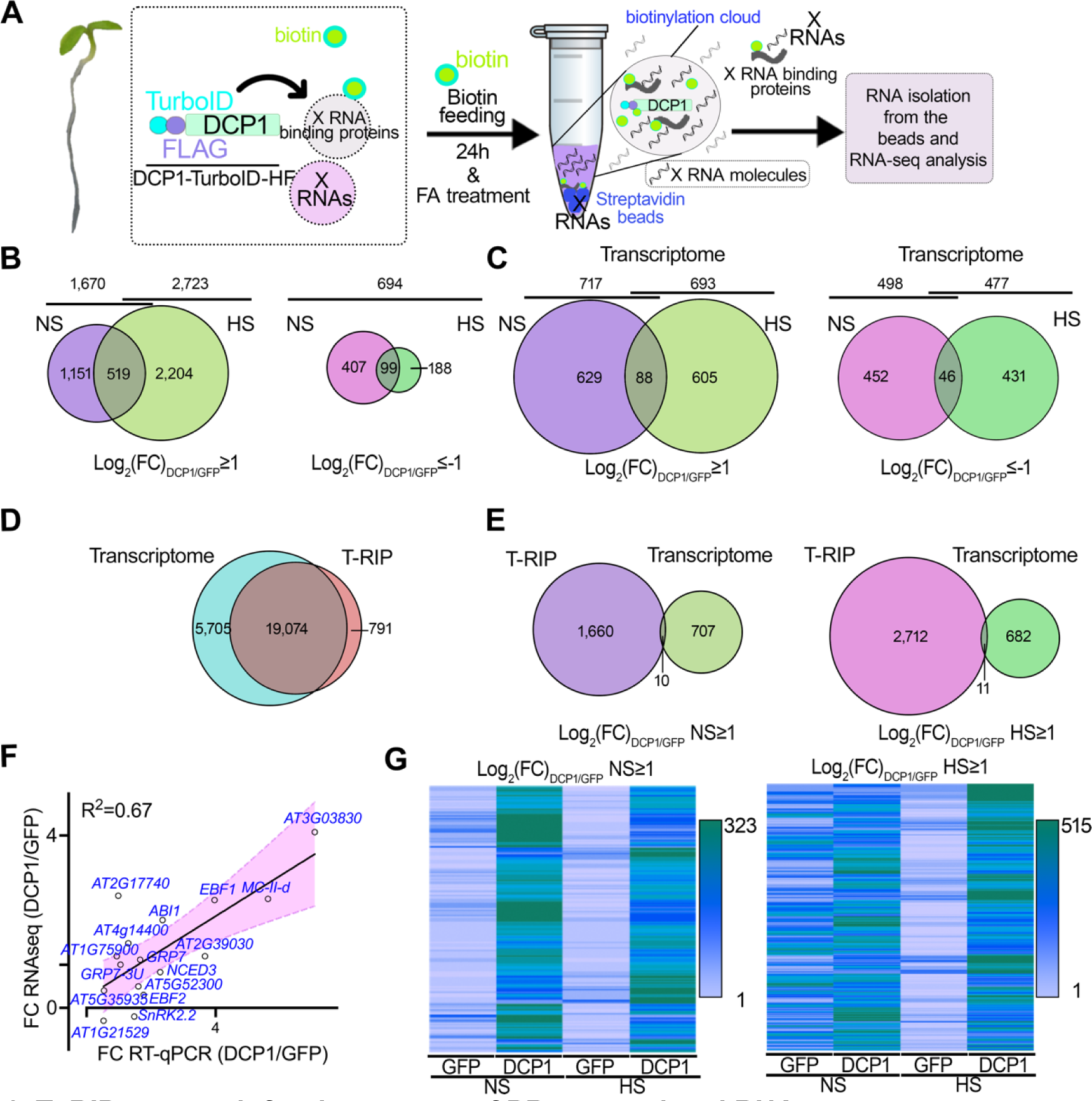
T–RIP approach for the capture of PBs–associated RNAs. **A.** Pipeline for T–RIP: 1. Plants were submerged in 50 μM biotin for 1 min under vacuum. The biotin solution was removed, and plants were placed back in the growth chamber. After 24 h, plants were left at the mock condition denoted as “non–stress” (NS) or treated with “heat stress” (HS; 22°C to 37°C for 2 h). 2. 1% (v/v) formaldehyde was used as an RNA-protein crosslinker (fed to tissues similarly to biotin). Formaldehyde was then quenched by glycine. 3. Proteins/RNAs were extracted as described for the APEAL approach (Liu et al., 2023); the extraction buffer was supplemented though with an RNase inhibitor. The supernatants were loaded on PD–10 columns to deplete excess biotin and were incubated with streptavidin–coated beads. 4. Protease treatments were used to elute RNP complexes from the beads, and RNA was extracted. All the T–RIP samples were subjected to DNase I treatment and ribosomal RNA depletion. **B.** Venn diagrams showing RNA identified for T–RIP (left; log_2_FC≥1) and those with a high probability to be excluded from PBs (right; log_2_FC≤–1; 506 RNAs in NS and 287 RNAs in HS). **C.** Venn diagrams showing genes identified for the transcriptome (total RNA–Seq) comparing the DCP1/GFP expressing lines (left, log_2_FC≥1; right, log_2_FC≤–1). Note that similar quantitative and small differences were found between samples regarding differential expression. **D.** Venn diagrams showing the overlap between unfiltered RNAs identified for the Transcriptome (from GFP and DCP1 samples) and T-RIP. **E.** Venn diagram comparing the overlap between the filtered enriched RNAs in NS (log_2_FC≥1) of the T-RIP dataset and of the corresponding transcriptome dataset in NS (left) and HS (right). Note: For aesthetic reasons and readability Venn diagrams in panels **B-E** are not proportional throughout and the analogies are kept only between the two conditions compared each time. **F.** Correlation between RNA levels determined by RNA–Seq or RT–qPCR for the indicated genes. *ACTIN7* and *PP2A* were used for normalization. Data are from three independent experiments with two technical duplicates (n=2 assays). The fitted line is shown along with the confidence intervals as shaded area (95%, deviation from zero *p*=0.0008), with a fitted equation Y = 0.4686*X + 0.2534. FC, fold change between DCP1/GFP samples. **G.** Heat maps showing sample gene clusters of T-RIP. RNAs were filtered either using a log_2_FC≥1 for genes enriched in DCP1 NS (left) or HS (right) from GFP and DCP1 samples. The color-coding bars indicate the number of RNAs.

Previously, using the *DCP1–TurboID–HF* lines, we achieved capturing the proteome of PBs in a tandem approach called “APEAL” [for Affinity Purification with Proximity– dEpendent LigAtion steps; (Liu et al., 2023)]. For T–RIP, we refined the APEAL using seedlings grown under normal conditions and transferred to non–stress (NS, 22°C, equals to normal) or HS (37°C) conditions for 2 h. We justify the use of HS considering that it leads to an increase mainly in the size of PBs and induces changes in PBs proteome (Liu et al., 2023). Seedlings were submerged in biotin and after 24 h RNAs were fixed to proteins using formaldehyde (**Fig. 1A**, details in Methods). Streptavidin– coated beads were used to enrich and purify PBs–associated RNPs and the corresponding RNA was extracted and subjected to RNA–Seq.

Due to the HS–induced changes in PBs proteome and considering that many proteins in PBs are RNA–binders, we speculated that these could be followed by consequent changes in PBs RNA composition. In accordance, although HS led to a decrease of the total protein content in PBs (from 16% to 6% of the proteins in APEAL), it induced the accumulation of RNA–binding proteins (**Supplemental File 1;** >10%, four-fold higher than the NS). We did RNA–Seq on NS and HS samples and considered only high– confidence enrichments (Fold enrichment (“FC”) log_2_FC_DCP1/GFP_≥1 or <-1, i.e., enrichment/depletion when comparing DCP1 to GFP samples; *n*=2, false–discovery rate [FDR]=0.05, **Supplemental File 2**). Multiplexed libraries averaged 20 million reads, mapping to the *Arabidopsis* genome >90%. T–RIP captured 19,865 RNAs of which 1,670 (8.4%) and 2,723 (13.7%) were enriched in DCP1 samples in NS and HS conditions, respectively (**Fig. 1B**). We found that 519 RNAs were common among NS and HS representing those that are likely always associated with PBs (**Supplemental File 3** for gene names). T–RIP DCP1 datasets had more RNAs enriched compared to GFP; only 694 RNAs were enriched in T–RIP GFP datasets (**Fig. 1B**, log_2_FC_DCP1/GFP_<-1; i.e., 3.49% from HS+NS). At the transcriptome level, however, HS induced similar quantitative changes in RNA levels for both NS and HS for up– and downregulated genes; ca. 500 RNAs were differentially expressed between DCP1 and GFP–TurboID in NS and HS (**Fig. 1C**). We could not identify in T–RIP dataset 5,705 RNAs (unfiltered, all RNAs from both NS and HS and DCP1 compared to GFP samples) that could be identified for the transcriptome, while 791 were exclusively identified for T–RIP (**Fig. 1D**). T–RIP dataset enriched RNAs were not significantly enriched in the total RNA sequencing dataset from the same samples using the same enrichment criteria; only 10 and 11 RNAs were enriched in both transcriptome and T–RIP for NS and HS, respectively (**Fig. 1E**). Hence, enriched RNAs in T–RIP dataset did not necessarily follow the corresponding levels of total RNA. In turn, among transcripts with a high enrichment in T–RIP were RNAs with a low read coverage in the total RNA (or absent). As mentioned above, there was little overlap between enriched RNAs in NS and HS (11% or 519 RNA), validating our hypothesis that HS not only remodels the PBs’ proteome but also their RNAs. To further validate T–RIP, we used RT–qPCR on T–RIP samples using 16 RNAs with various enrichment levels (log_2_FC –0.3 up to 4.1). The RNA levels determined by RT– qPCR correlated relatively well with the corresponding FC obtained in T–RIP (**Fig. 1F**, R^2^ = 0.67; **Supplemental File 4**, genes used). We also confirmed the presence in our T–RIP datasets of the few known RNA molecules localizing at PBs, *EBF1* (*EIN3 BINDING F–BOX PROTEIN1*), and *EBF2* [(Li et al., 2015; Merchante et al., 2015); **Fig. 1F** and **Supplemental File 4**]. Furthermore, T–RIP enriched RNAs in DCP1 samples showed certain enrichment clusters in heat maps that were not observed in GFP controls (**Fig. 1G**).

As a cautionary note here, we used T–RIP lines overexpressing DCP1 (*DCP1* mRNA increased by ∼30%, (Liu et al., 2023)). We thus compared whether DCP1 expressed under a native promoter (*DCP1pro*) could also lead to the enrichment of selected RNAs. We fixed with formaldehyde seedlings expressing DCP1–GFP under the *35Spro* or *DCP1pro* at NS (similar conditions used for T–RIP). We further purified enriched RNAs following a refined approach for GFP–trapping (with GFP–beads adjusting formaldehyde fixation). We selected as a readout RT–qPCR of five genes enriched and three depleted (with high confidence) from T–RIP. We observed similar enrichment patterns between *35Spro* and *DCP1pro* lines when normalized to a line expressing GFP (**Supplemental Fig. 2**). Yet, these enrichments were moderate when compared to those obtained using T–RIP (e.g., for EBF2 2 for T-RIP vs. 1.3 for GFP-trap), consistent with the shorter radius of formaldehyde fixation than the biotinylation radius of TurboID; we have suggested that TurboID can indeed retain interactions not identified by formaldehyde fixations or simple affinity purifications (Liu et al., 2023). Noteworthy, is the high enrichment of the actin–related *ADF9*, consistent with our previous results on the SCAR–WAVE–DCP1 axis. Furthermore, through this approach, we likely identify RNAs associated with the DCP1–DCP2 complex that can exist outside of the condensed phase. We thus acknowledge that overall RNA enrichments in PBs could differ to some extent from the ones we define here. Collectively, we propose that T–RIP can capture at least a part of RNAs localizing in PBs; we refer hereafter to these RNAs as “PBs–enriched RNAs”.

We classified the PBs–enriched RNAs according to their Gene Ontology (GO, Biological Process) using PANTHER (protein analysis through evolutionary relationships; cutoff *p*<0.05 for the –log_2_FDR, **Supplemental Files 3** and **5**, GO redundant terms are present). Most RNAs in NS and HS were classified in four GO subnetworks: 1. membrane/cell wall, 2. nucleic acid metabolism organization/RNA localization or transport, 3. assembly of macromolecular complexes, and 4. plant metabolism/development [e.g., regulation of hormonal responses, auxin, and ethylene; **Fig. 2A**)]. GOs within these four subnetworks mostly appearing in NS conditions, related to wounding/regeneration [**Fig. 2A**; indicated with blue boxes, i.e., defense response, secondary metabolism, glucosinolates–cell wall (FDR = 3.97e^−3^), callose deposition (FDR = 3.46e^−3^), and actin organization (FDR = 4.58e^−2^)]. On the contrary, GOs such as light signaling were observed in HS conditions [**Fig. 2A**; shade avoidance and response to light stimulus (FDRs = 3.12e^−2^ and 4.59e^−4^)]. The involvement of PBs in the regulation of light signaling has been previously described (Jang et al., 2019) and is consistent with our results here. We also observed common GOs between NS and HS, e.g., of hormonal responses (indicated with magenta boxes, FDR, NS = 5.15e^−5^, HS = 6.04e^−5^) and actin (**Fig. 2A**). The PBs–depleted RNAs fit into two subnetworks related to the housekeeping processes of nucleic acid and plant metabolism/development (**Fig. 2B**). The enrichment distribution of RNAs in PBs is shown in **Fig. 2C**, where we highlight enriched RNAs from the four subnetworks. Altogether, we conclude that PBs–enriched RNAs can fit into a small number of conditional subnetworks.

**Figure 2.**
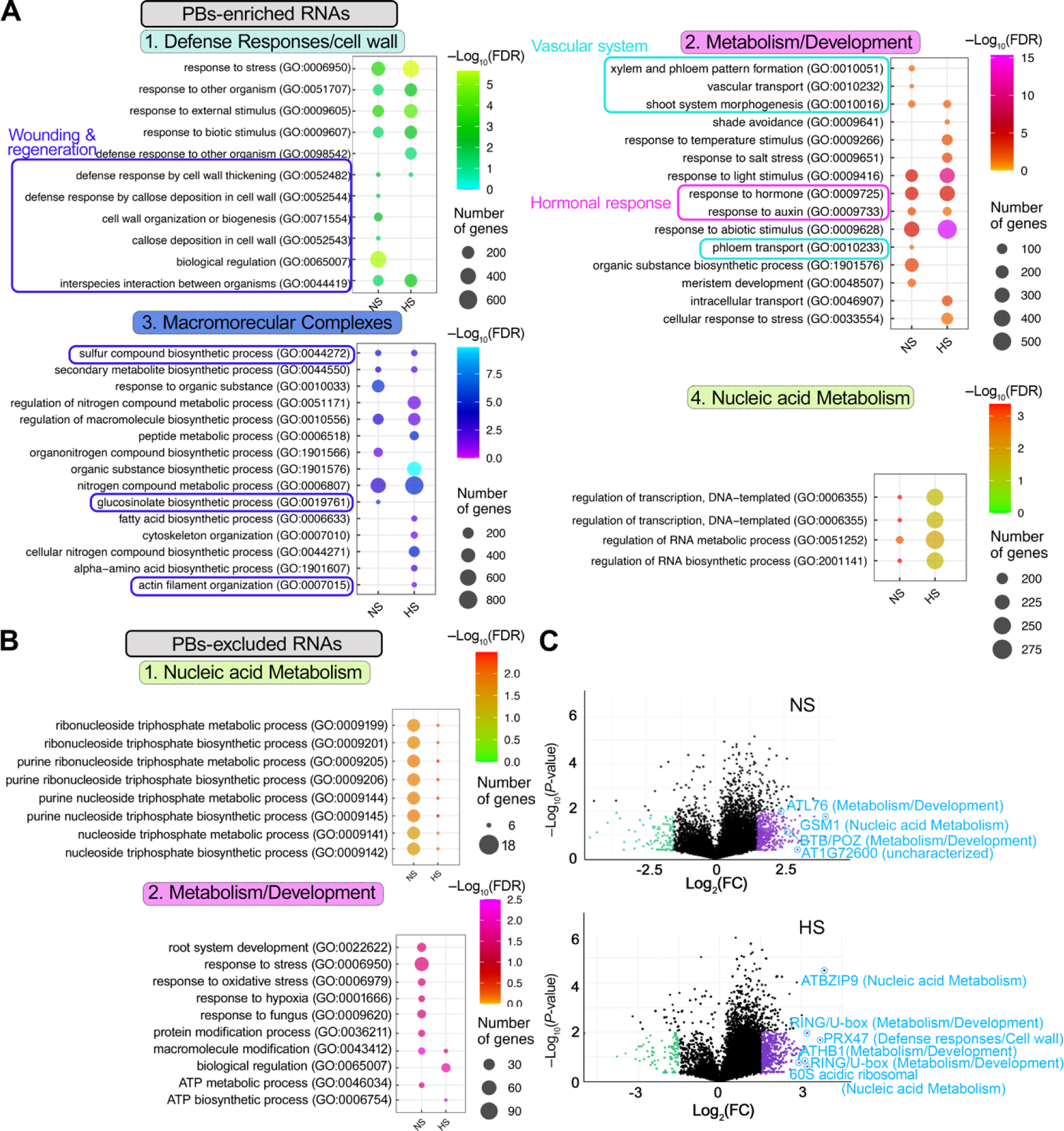
Subnetworks of RNAs enriched in PBs. **A.** GOs of PBs-enriched RNAs fitting into four subnetworks. Boxes annotate clusters: wounding and regeneration (blue), vascular system (light blue), and hormonal responses (magenta). FDR, false discovery rate. **B.** GOs of PBs-excluded RNAs fitting into two subnetworks. **C.** PB-enriched/depleted RNAs (log_2_FC_DCP1/GFP_) in NS (upper) or HS (lower), visualized by volcano plots. RNAs from the four subnetworks defined in **A** are indicated.

### RNAs with shorter coding regions tend to localize in PBs

Thus far, we have defined the PBs–enriched RNAs under two conditions. Our next question was whether these RNAs have specific features under these conditions that promote their recruitment to PBs. In animals, RNA size seems to modulate PBs localization (Wang et al., 2018). We thus first established a pipeline of comparison between PBs–enriched/depleted RNAs and total transcriptome for 3’– or 5’–UTR, coding sequences (CDS) lengths, and GC% content. By examining and analyzing these parameters in an unbiased approach, we found a negative correlation between CDS length and PBs association (r = ∼–0.25 in NS and HS, while r >0.2 for total; **Supplemental Fig. 3A–I**). Hence, smaller RNAs tend to localize in PBs under NS and HS conditions.

We next queried secondary structures’ free energy (folding energies) of PBs–enriched RNAs following the same pipeline described above. In yeast and human cells, the 5’– UTRs secondary structure regulates mRNA localization (Ringner and Krogh, 2005). Accordingly, considering that the size of UTRs is not affecting localization in PBs, we opted for using a normalized metric, the “minimum free energy (MFE) density” that discounts size and works for RNA >40 nt (Trotta, 2014). This metric accounts for local free energy, which could be important for RNA functions in PBs. High structural stability is compatible with specific functions, for example, the necessity of RNAs to maintain a stem–loop structure for recognition by RNA–binding proteins (Ha and Kim, 2014). Indeed, MFE density is likely more adequate for comparisons in PBs–enriched RNAs, as it correlated well with GC% but not, as expected, with length (**Supplemental Fig. 4A–D**). We observed that increased stability of 5’–UTR (lower MFE density), was associated well with increased half–life [as defined in (Sorenson et al., 2018)]; yet PBs– depleted RNAs had increased 5’–UTR stability (**Supplemental Fig. 5**). This result suggests that PBs may help in the stabilization of RNAs, stabilizing RNAs with loose UTRs. Accordingly, from the same datasets, we observed that HS was associated with higher stability than NS for most RNAs, while the small size of CDS in NS was associated with reduced RNA stability. We did not find though a correlation between enrichment level and MFE density (r<0.01, in NS and HS). During these analyses, we also observed that PBs-enriched UTRs did not show the composition bias for increased adenine and guanidine which correlates with decay [**Supplemental Fig. 6**, defined in (Sorenson et al., 2018)]. As PBs can compete with polysomes for RNAs, we also tested whether PBs–enriched RNAs have codons that could halt translation and shuttle RNAs to PBs for decay. Using the Relative Codon Bias Strength (RCBS) index to calculate the difference between the observed versus the expected codon frequency (Sahoo et al., 2019; Parvathy et al., 2022), we observed a marginally increased RCBS in HS (**Supplemental Fig. 7A–C**). Overall, these results suggest that the small size of RNA and a looser 5’–UTR structure associate with localization to PBs.

### Modification of N^6^–methyladenosine is not a prerequisite for PBs–RNA enrichment

Thus far, our results predict a moderate link between RNA primary and secondary structure with PBs localization. We thus further asked whether PBs–enriched RNAs show modifications that could modulate their localization with PBs. We focused here on N^6^–methyladenosine (m^6^A), owing to its high abundance (25% of all RNAs are modified) and the ability to modulate PBs dynamics in non–plants (Zaccara and Jaffrey, 2020). To check whether PBs RNAs are m^6^A–enriched, we compared PBs–enriched/depleted RNAs to the dataset described in (Parker et al., 2020). In this work, to extract information about m^6^A–enriched RNAs, authors compared RNA–seq reads of the *VIRILIZER* (*vir–1*) mutant that represents a conserved m^6^A component of the writer complex, to a corresponding complementation line (VIR–GFP, denoted as “virC”). This comparison confirmed that PBs–enriched RNAs can bear m^6^A–modifications, mainly during HS (**Supplemental Fig. 8**).

To experimentally verify this link we used a mutant allele of the m^6^A writer FIP37 [FKBP12 Interacting protein 37; *fip37*–*4* (Shen et al., 2016)], comprising the writer complex, and showing an ∼85% reduction in m^6^A. We introduced *DCP1pro:DCP1–GFP* in the *fip37–4* background by crossing. The *DCP1–GFP fip37–4* mutant did not differ phenotypically from *fip37–4* and showed similar to wild type levels of *DCP1* expression (**Fig. 3A, B**). We observed reduced stability of PBs in *DCP1–GFP fip37–4* as determined by a short treatment with cycloheximide that is known to disassemble PBs (Gutierrez-Beltran et al., 2015), and an increase in PBs sizes/numbers and fluorescence recovery after photobleaching [FRAP; **Fig. 3C**]. *DCP1–GFP fip37–4* showed an unexpected increase in size/number of PBs (**Fig. 3C**). Rapid rates in FRAP are diagnostic of high liquidity and likely reduced stability of condensates (Alberti et al., 2019). Furthermore, treatments with 1,6–hexanediol that dissolves liquid condensates while leaving the more solid ones intact (Alberti et al., 2019), showed that *DCP1–GFP fip37–4* PBs could be dissolved similarly to wild type ones (**Fig. 3D**). Hence, our results suggest that m^6^A–modification on RNAs may modulate stability but not the formation of PBs.

**Figure 3.**
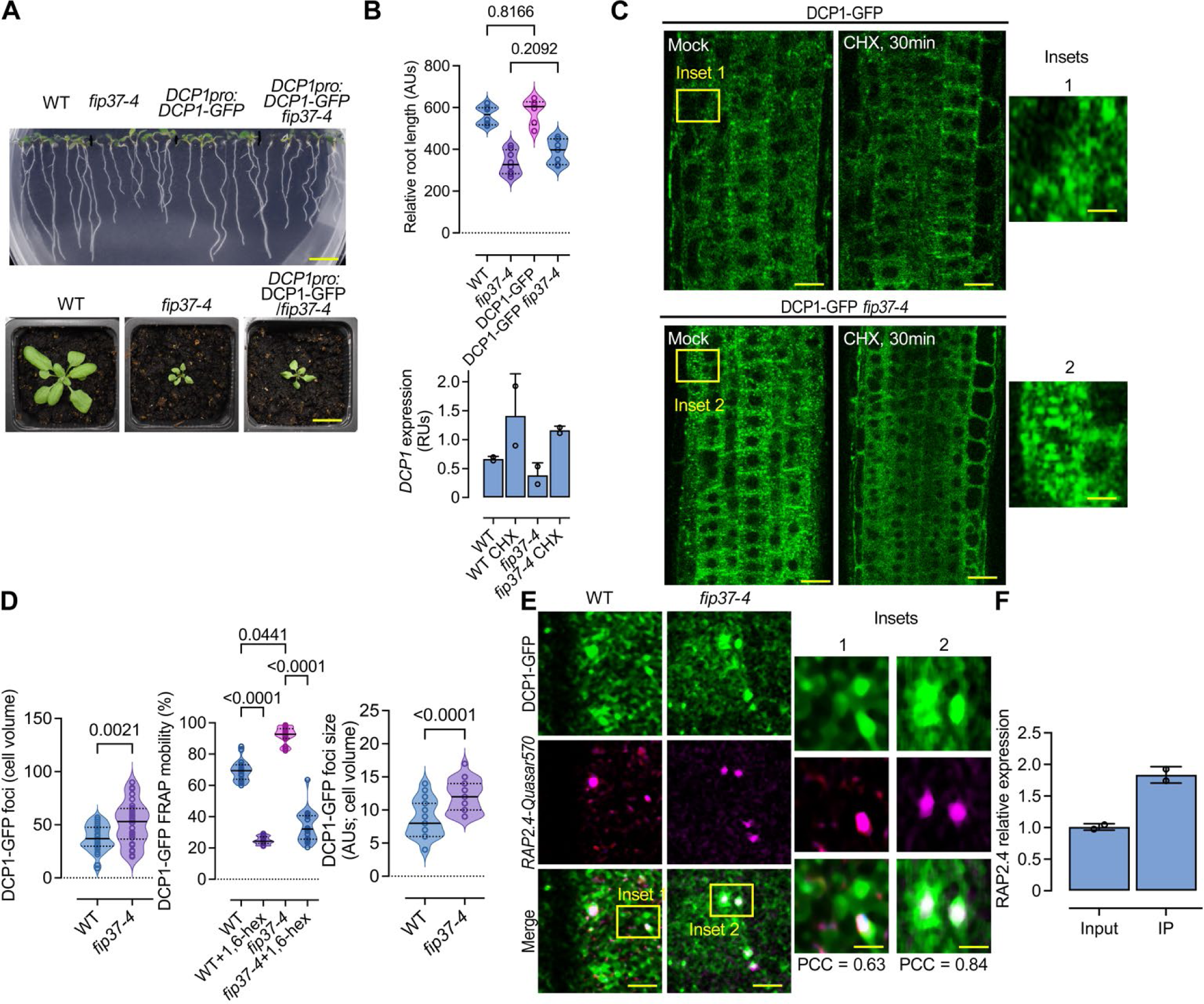
The m^6^A writer FIP37 controls stability but not the formation of PBs. **A.** Phenotype of *fip37-4* or *fip37-4 DCP1pro:DCP1-GFP* seedlings (7 days after germination (DAG)). Lower: corresponding phenotypes of rosettes. Length bars, 1 cm. **B.** Relative root length of wild type (WT), *fip37-4* or *fip37-4 DCP1pro:DCP1-GFP* seedlings. Data are means of three independent experiments with two technical replicates (n=2 with 10 roots from 7 DAG seedlings), and significance was determined by ordinary one–way ANOVA. Lower: RT-qPCR of DCP1 expression in WT or *fip37-4 DCP1pro:DCP1-GFP* seedlings (7 DAG), in the presence or absence of cycloheximide (CHX, 30 μM, 30 min). *TUA4* was used for normalization. Data are from two independent experiments with two technical duplicates (n=2 assays). AUs, arbitrary units; RUs, relative units (normalized to *TUA4*). **C.** Micrographs from meristematic epidermis root cells 5 DAG expressing *DCP1pro:DCP1–GFP* in WT or *fip37-4* in the presence or absence of cycloheximide (CHX) (30 μM, 30 min). The experiment was replicated three times. Scale bars, 10 μm. AUs, arbitrary units. **D.** Quantification of the number of DCP1-GFP foci per cell (cell volume in μm^3^), corresponding FRAP mobility (%; corresponding to the initial signal recovery), and diameter (size) of PBs in WT or *fip37-4*. For FRAP mobility, seedlings were treated with 1,6-hexanediol (“hex”), to dissolve PBs showing liquidity. Data are means of three independent experiments with two technical replicates each (n=2 with 6-10 roots from 5 DAG); significance was determined by ordinary one–way ANOVA. AUs, arbitrary units. **E.** Micrographs from meristematic epidermis root cells 7 DAG showing smFISH detection of *RAP2.4-Quasar570* mRNA (in magenta) expressing *DCP1pro:DCP1-GFP* in WT or *fip37-4*. Two independent experiments with (n=6-10 cells randomly picked) showed similar results. Scale bars, 2 μm (insets, 0.2 μm). **F.** Immunoprecipitation (IP)-RT-qPCR of m^6^A-modified *RAP2.4*. Data are means of two independent experiments with two technical replicates each (n=2 IP assays from 5 DAG seedlings). Data were normalized against tubulin which is also m^6^A-modified (Parker et al., 2020). The technical control for the immunoprecipitation was the one suggested by the manufacturer (Materials, m6A modified Control RNA (*Gaussia Luciferase*)).

We further queried whether the PBs in the *DCP1–GFP fip37–4* showed reduced localization of PBs–enriched RNAs, for example, the *WOUND INDUCED DEDIFFERENTIATION 1/RELATED TO AP2 4* [*RAP2.4*, log_2_(FC)_NS,HS_=1.586/1.943] (**Fig. 3E**). We justify this selection according to RAP2.4 relevance to hormonal responses and regeneration, processes followed up later in this paper. We thus adopted and refined a single–molecule fluorescence *in situ* hybridization (smFISH) approach in root meristem cells enabling the observation of intact tissues (Zhao et al., 2023). For smFISH, we used labeled probes with the fluorescence dye Quasar 570 spanning the whole CDS (see also Materials). We observed a slight increase of the *RAP2.4* signal in *DCP1–GFP fip37–4* PBs when compared to the corresponding wild type (**Fig. 3E**). Furthermore, *RAP2.4* is m^6^A modified under our conditions, as shown using m^6^A-immunoprecipitation (IP) combined with RT-qPCR (**Fig. 3F**). These results suggest that m^6^A–modification levels may not necessarily affect RNA localization to PBs. Although m^6^A–modifications may not be necessary for the recruitment of RNAs to PBs, they could modulate PBs’ stability. The exact mechanism of this modulation merits further investigation and likely involves multiple regulatory pathways affecting the stability and interactions of RNAs.

### PBs size inversely correlates with their RNA degradation capacity

Because RNA decay factors accumulate in PBs, it was proposed that PBs’ function is that of RNA decay (Luo et al., 2018). On the other hand, our four observations so far suggest that PBs may also store RNAs in plants: 1. m^6^A–RNAs which are likely enriched in PBs are generally more stable [for example, (Song et al., 2023)]. Furthermore, PBs–enriched UTRs were not rich in adenine and guanidine that associate with decay [**Supplemental Fig. 6** and (Sorenson et al., 2018)]; 2. During HS, when PBs increase in size, we found more RNAs in PBs than in NS conditions (**Fig. 1B**). Furthermore, smaller RNAs which are mainly associated with PBs, show reduced decay in HS. This finding is consistent with observations in yeast where PBs store RNAs upon glucose starvation for later translation (Wang et al., 2018); 3. The RNA–seq read distributions, calculated using “metagene plots” that provide profiles of average coverage of PBs–enriched RNAs showed a slight reduction towards the 3’–UTR consistent with the association of the de–adenylation machinery with PBs (**Supplemental Fig. 9**). Accordingly, in non–plants, de–adenylation has been proposed as a prerequisite for RNA association with PBs, prior to decapping (Chen and Shyu, 2013). On the other hand, we did not find the expected reduction of 5’–UTR reads (especially during HS), as the decapping would trim 5’–UTRs through XRN4 or other exonucleases (Goeres et al., 2007). In accordance, we did not find a correlation between the levels of our previously obtained decapping dataset (from NS and HS) with the enrichment level of PBs–enriched RNAs; we should note here that we found a moderate negative correlation with FC of PBs-excluded RNAs confirming their reduced decapping [**Supplemental Fig. 10A**; dataset from (Gutierrez-Beltran et al., 2015)]; 4. Finally, when comparing the degradation rates for total transcriptome defined in Sorenson et al., (2018) (for both wild type and *varicose* mutant) with PBs–enriched RNAs with those excluded we found that in HS, when PBs increase in size, RNAs were rather more stable (**Supplemental Fig. 10B**).

We thus asked when could PBs function in RNA decay by measuring the effect of PBs numbers/sizes in RNA half–lives. To address this question, we first used 5–d–old seedlings treated with a transcription inhibitor (cordycepin) to block *de novo* RNA synthesis and collected samples across a 240 min time course (Sorenson et al., 2018). We compared RNA decay between wild type and *dcp1–3* with fewer PBs or the SCAR/WAVE complex mutant *scar1234* with more and larger PBs [**Supplemental Fig. 11** and (Liu et al., 2023)]. We followed the degradation rates of three PBs–enriched RNAs using RT–qPCR according to their relevance to hormonal responses and regeneration, processes followed up later in this paper, *RAP2.4*, *EBF2* [log_2_(FC)_NS,HS_= 1.015/1.205], and the *RELATED TO AP2.4* [*RAP2.4D/WIND2*; log_2_(FC)_NS,HS_ = 1.426/1.550; class 2 of CDS length, 800–1,600 close to the expected 1.2]. When compared to the wild type, the *scar1234* showed increased, while *dcp1–3* reduced degradation rates (mainly for *WIND2*); the ranges of decay defined were similar to the ones described previously [(**Fig. 4A, B**; (Sorenson et al., 2018)].

**Figure 4.**
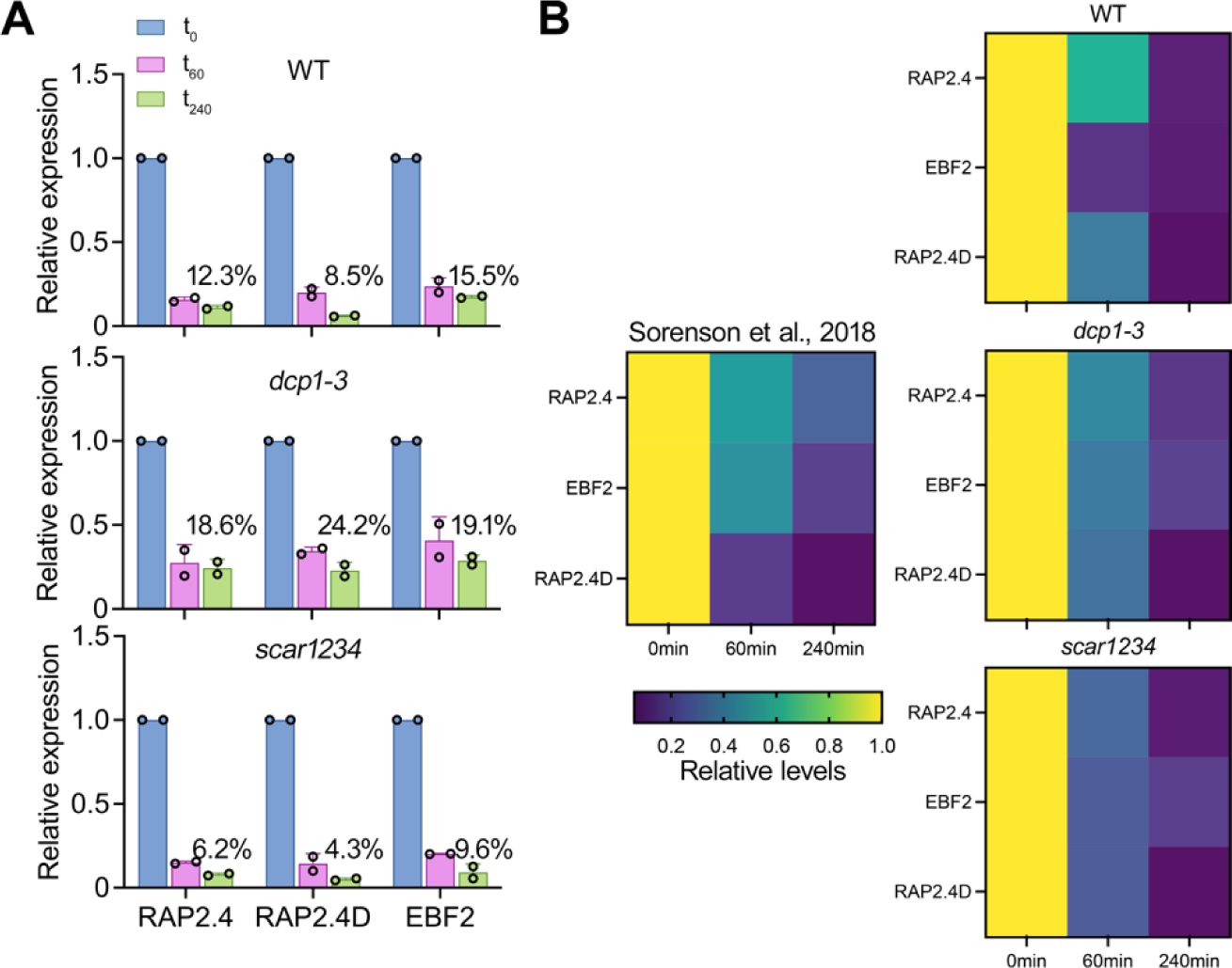
Decay rates modulation by the number and size of PBs of three PBs-enriched RNAs. **A.** RT-qPCR determination of RNA decay levels for *RAP2.4*, *EBF2,* and *RAP2.4D*, in wild type (WT), *dcp1-3,* and *scar1234* 5 DAG and in three time points (0, 60, and 240 min) upon treatment with the transcription inhibitor cordycepin [1 mM, (Sorenson et al., 2018)]. The percentages show the levels of RNAs at 240 min compared to 0 min. Data are means of two independent experiments with two technical replicates each (n=2 roots from 5 DAG seedlings). **B.** Heat maps showing RNA decay levels for *RAP2.4*, *EBF2,* and *RAP2.4D* were obtained from (Sorenson et al., 2018) and compared to the defined herein decay rates in WT, *dcp1-3,* and *scar1234* lines defined by RT-qPCR.

As *scar1234* showed both increased numbers and sizes of PBs, we asked which of the two parameters could contribute to the increase of RNA decay. As the association of DCP1 and DCP2 correlates well with active RNA–degradation (Tibble et al., 2021; Liu et al., 2023), we aimed at using DCP1–DCP2 interaction as a proxy for decapping/decay.

To achieve this aim, we established a quantitative 3D proximity ligation assay (PLAs; (Teale et al., 2021)). PLA uses complementary oligonucleotides fused to antibodies to determine the frequency with which proteins of interest find themselves nearby (**Fig. 5A**). PLA allows *in situ* detection of endogenous protein interactions with high sensitivity and single–molecule resolution (at distances < 40 nm). Typically, two primary antibodies against epitope tags are used to detect two unique protein targets. A pair of oligonucleotides–labeled secondary antibodies (PLA probes) bind to the primary antibodies. Next, hybridizing connector oligos join the PLA probes only if they are near each other, and ligase forms a closed, circle DNA template that is required for rolling– circle amplification (RCA). The PLA probe then acts as a primer for a DNA polymerase, generating concatemeric sequences during RCA. This RCA step allows up to a 1,000– fold signal amplification that is still tethered to the PLA probe, allowing the detection of even very transient interactions. Last, labeled oligos hybridize to the amplicon, which is then visualized and quantified as discrete spots (PLA signals). Contrary to our expectations, we discovered that DCP1–DCP2 interacted well in submicroscopic PBs of root epidermal meristematic cells (i.e., on the limit of or below detection in our super– resolution confocal setting, 120 nm). We assume that these PBs could even represent “RNA–singletons”, that is single or few RNA molecules with the decapping complex attached (Mateu-Regue et al., 2019). By correlating DCP1–DCP2 interaction (PLA signals) with the size of PBs through regression analyses, we found a strong negative correlation (**Fig. 5B**; R^2^=–0.6). As DCP1–DCP2 colocalized in most PBs (**Fig. 5B**; ∼80%), this result suggests that DCP1–DCP2 usually dissociate in larger PBs (>∼400 nm).

**Figure 5.**
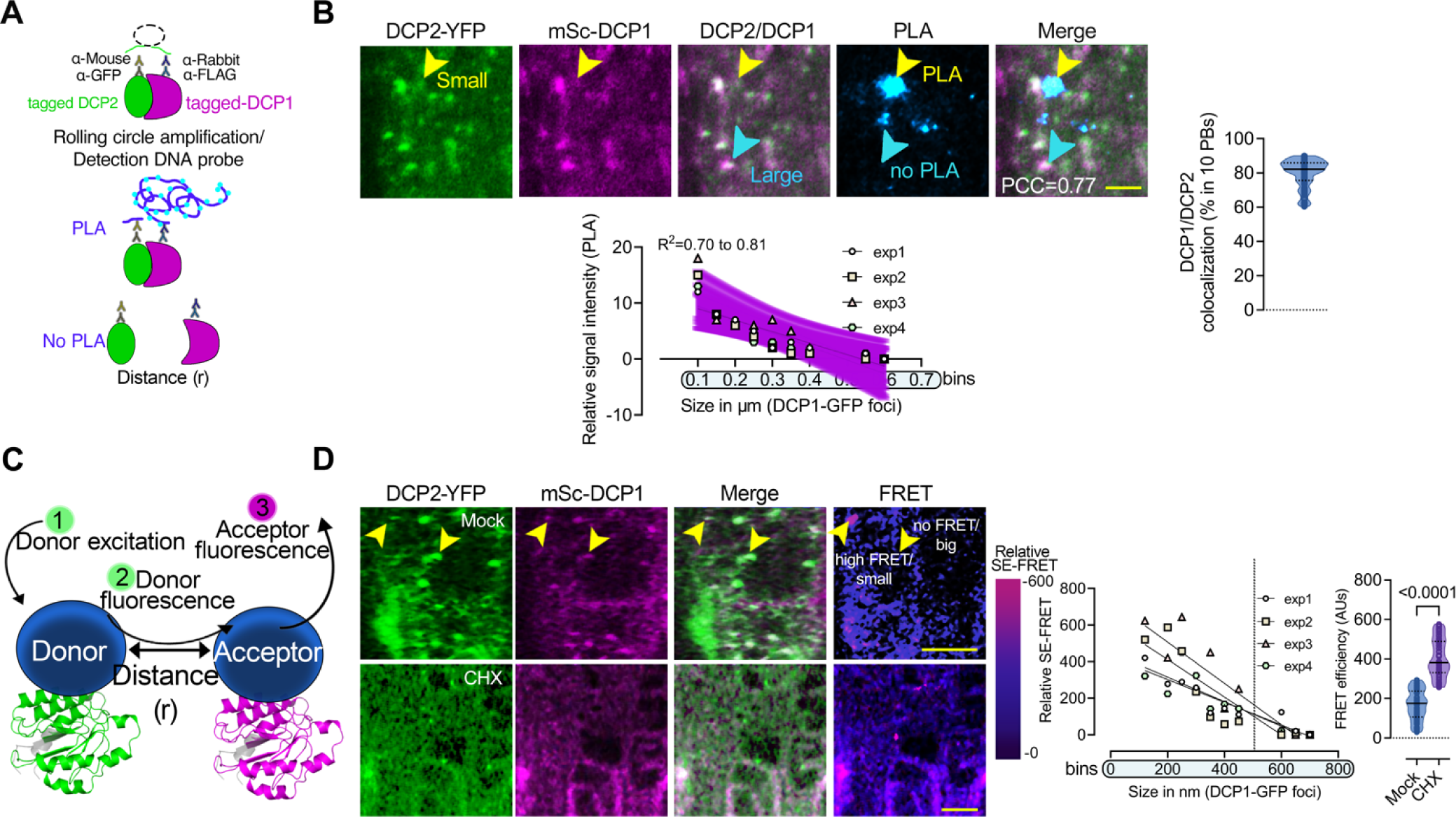
Correlation between DCP1/DCP2 interaction with the size of PBs in root cells. **A.** Illustration of the PLA approach principle (see details in the main text). **B.** PLA signal produced by α–GFP/α–FLAG of a line co–expressing *RPS5apro:HF–mScarlet–DCP1* (mSc-DCP1) and *35Spro:DCP2–YFP*. The “spots” (violet “PLA”) do not connote physiologically relevant puncta (e.g., condensates). The co–localization of PLA spots with DCP1-DCP2 signals is also shown [noted as Pearson correlation coefficient (PCC)]. Scale bars, 5 μm. Lower: correlation between the size of PBs (DCP1-GFP foci) and PLA spot number (DCP1/DCP2 interaction), in 100 nm PBs size bins. Data are means of four independent experiments with two technical replicates each (n=2 roots with 5 meristematic epidermal cells 5-7 DAG each). The simple regression analysis includes data from the four independent experiments denoted as “exp”. All fitted lines are shown along with the overlaid confidence intervals (purple stripe; 95%, deviation from zero *p*=0.0009–0.0049), with a fitted equation Y = –20.54*X + 11.06). Non–linear regression through one-phase decay predicted relevant models here. Right: colocalization of DCP1/DCP2 per ten PBs. Data are means of four independent experiments with two technical replicates each (n=2 roots with 5 randomly selected meristematic epidermal cells 5-7 DAG). **C.** Sensitized emission FRET principle (SE-FRET), where the emission spectrum of the donor (1) overlaps with the excitation spectrum of the acceptor (2), and if the distance (r) between the two molecules is sufficient, energy is transferred (3). **D.** Micrographs of root meristematic epidermal cells showing SE–FRET efficiency between mScarlet– DCP1 and DCP2–YFP in the absence or presence of cycloheximide (CHX, 10 μM 20 min). The arrowheads denote small (at the detection limit) or big PBs (upper micrograph). Scale bars, 10 μm. Right: correlation between PBs size and SE–FRET efficiencies in 50 nm PBs size bins and SE-FRET efficiency in the presence or absence of CHX. Data are means of four independent experiments with two technical replicates each (n=2 roots in each replicate with 10 randomly picked meristematic epidermal cells 5-7 DAG). The simple regression analysis includes data from four independent experiments denoted as “exp” (n=16–33 cells each). All fitted lines are shown.

To discount the possibility that PLA antibodies could not identify some DCP1/DCP2 molecules due, e.g., to stereochemical hindrance in the crowded and viscous environment of PBs, we refined a live cell imaging approach. We thus used the stoichiometry–sensitive type of Försters resonance energy transfer called “sensitized emission” (**Fig. 5C**, SE–FRET). FRET is a non–radiative interaction between two molecules that happens at distances in the range of a few nm. In SE–FRET, the sensitized emission (i.e., the acceptor fluorescence resulting from energy transfer from excited donor molecules) from separately acquired donor and acceptor images is calculated, following leak–through corrections (van Rheenen et al., 2004). The SE– FRET approach allows FRET measurements on moving condensates (e.g., PBs) and could in principle provide stoichiometric information about DCP1–DCP2 interactions within a shorter range than PLA (i.e., 10 nm). Using SE–FRET, we could confirm PLA results (**Fig. 5D**). Furthermore, using a short treatment with cycloheximide to partially reduce the size and dissolve PBs we observed a transient increase of DCP1–DCP2 interaction (**Fig. 5D**, CHX). Together, PLA and SE–FRET results implied that RNA decay is mainly executed by small PBs (or in diffused decapping complexes).

To validate this suggestion, we directly queried RNA fate in PBs using smFISH. We performed quantitative smFISH colocalizations between PBs labeled by the DCP1–GFP and three RNAs enriched in PBs with sets of Quasar 570 labeled mRNA probes (*EBF2*, *DFL1,* or *RAP2.4*; **Fig. 6A**). As a negative control, we used the PBs–depleted RNA *SERINE/THREONINE PROTEIN PHOSPHATASE 2A* [*PP2A;* log_2_(FC)_NS/HS_ =0.46/0.15; **Supplemental File 6** for enrichments of RNAs used for probes). We could confirm the colocalization between PBs and *EBF2*, *DFL1,* or *RAP2.4*, and the lack of colocalization with *PP2A* (**Fig. 6 Α, Β**). Using high–resolution imaging, we found a positive correlation between large PBs (>400 nm) and localization of the smFISH signal, with size up to a point (**Fig. 6C**). We also applied an ethylene precursor short treatment (10 μΜ 1– aminocyclopropane–1–carboxylic acid, ACC) for *EBF2*, as it can increase *EBF2* localization with PBs [**Fig. 6C**; see also **Supplemental Fig. 15** and (Li et al., 2015; Merchante et al., 2015)]. mRNAs, in general, showed a similar trend of accumulation in PBs with that observed using the above probes, as revealed using co–staining of the *RAP2.4* with Cy5–labeled oligoDT probes (**Fig. 6D**). Furthermore, treatments with cordycepin affected the accumulation of *RAP2.4* only in smaller PBs and had no effect for larger ones (**Fig. 6E**). Overall, we confirm *in vitro* observations and models for decay for non–plants (Tibble et al., 2021), showing that small PBs execute RNA decay *in vivo* (the corresponding model is presented in **Fig. 6F**).

**Figure 6.**
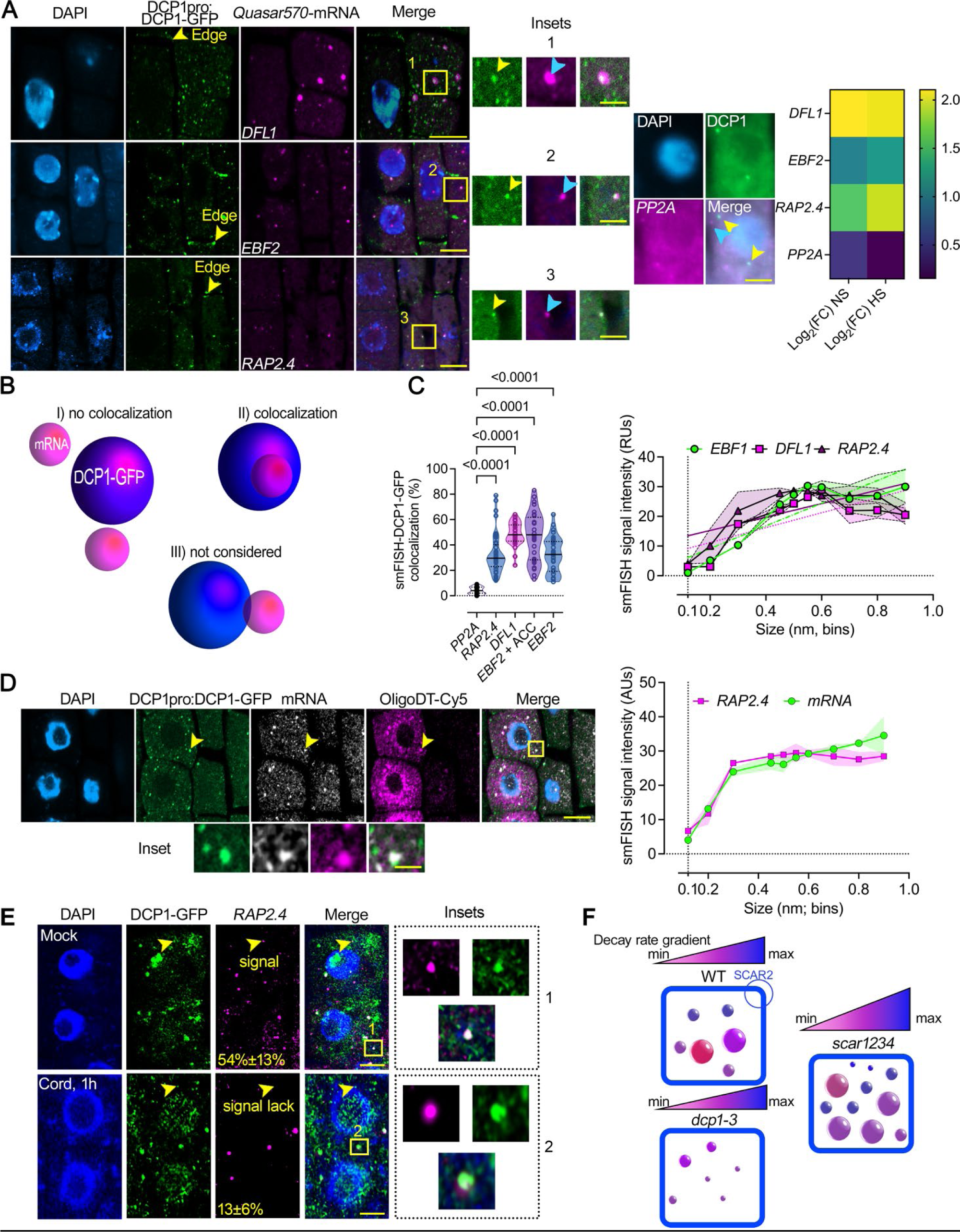
*In vivo* correlation between PBs size and RNA levels. A. Micrographs from smFISH for detection of *DFL1*, *EFB2*, RAP2.4, and *PP2A* mRNAs (in magenta) in root meristematic epidermal wild type (WT) cells 5 DAG expressing *DCP1pro:DCP1–GFP*. Insets: details of DCP1–GFP with each mRNA probe and with the negative control PP2a mRNA probe showing puncta colocalization (black box; arrows). Nuclei were stained with DAPI (violet). Right: DCP1-GFP/*PP2A* lack of colocalization and heat map showing enrichment levels for the four RNAs. The experiment was replicated three times (n=10 randomly selected cells). Scale bars, 2 μm (insets, 0.5 μm). **B.** Cartoon displaying the strategy for quantifications of colocalization between DCP1–GFP and mRNAs smFISH signal. Three major classes of colocalization are shown (I–III). Case III was not considered as co–localization since confocal microscopy may underestimate distances. **C.** Colocalization efficiency between mRNAs/DCP1–GFP (number of RNA smFISH puncta in PBs/total number of RNA puncta expressed as a percentage). Data are means of three independent experiments with two technical replicates each (n=2 roots with 10 meristematic epidermal cells 5 DAG), and significance was determined by ordinary one–way ANOVA. Eth, ethylene in the form of 10 μΜ ACC (see also **Supplemental Fig. 15**). Right: corresponding quantifications of the correlation between mRNAs/DCP1–GFP and PBs size in 100 nm PBs size bins. Data are means of three independent experiments with two technical replicates each (n=2 roots with 10 randomly selected meristematic epidermal cells 5 DAG each); significance for the violin plots was determined by ordinary one–way ANOVA. For the line plot, data are means±SD (SD: denoted as shaded area; N=3 biological replicates with n=10 randomly selected meristematic epidermal cells 5 DAG each). Fitted lines are also shown. **D.** Micrographs from *RAP2.4* smFISH signal (grey) counterstained with oligoDT-Cy5 in root meristematic epidermal wild type cells expressing *DCP1pro:DCP1–GFP*. Nuclei were stained with DAPI (violet). Arrowheads denote an example of the colocalization between *RAP2.4*, OligoDT, and DCP1-GFP, and the inset below denotes a detail of this colocalization. Right: quantification of the correlation between PBs size (DCP1-GFP foci) and *RAP2.4*/OligoDT signals in 100 nm PBs size bins. For the line plot, data are means±SD denoted as the shaded area (N=3 biological replicates with n=10 randomly selected meristematic epidermal cells 5 DAG each). Fitted lines are also shown. **E.** Micrographs from smFISH for detection of *RAP2.4* (in magenta) counterstained in the presence or absence of cordycepin (cord, 1 h), in root meristematic epidermal WT cells 5 DAG expressing *DCP1pro:DCP1–GFP*. Nuclei were stained with DAPI (violet). Arrowheads denote small PBs with or without smFISH signal for mock or cordycepin treatments. Percentages on micrographs indicate small PBs (∼0.2 μm), with smFISH signal ± SD. The two values were statistically different at *P*<0.05 (n=10 randomly selected meristematic epidermal cells; one-way ANOVA). Insets on the right, indicate large PBs with smFISH *RAP2.4* signal for mock and cordycepin treatments. The experiment was replicated three times (n=10 randomly selected cells). Scale bars, 5 μm (insets, 0.2 μm). **F.** Graphical representation PBs sizes in WT, *dcp1-3,* and *scar1234* meristematic, epidermal root cells. Above each model, there is a graphical representation of the decay rate. The *scar1234* contains a larger range of turnover. SCAR2 is the main protein responsible for retracting DCP1 from PBs and thus, their dissolution (Liu et al., 2023).

### An increase in protein disorder and RBPs in PBs correlates with RNA storability

We next aimed to determine PBs features that could associate with increased size and RNA storage. *In silico* analyses of PBs proteome through the PLAAC algorithm (Dosztanyi et al., 2005) obtained through APEAL (Liu et al., 2023), revealed that only during HS, when PBs increase mainly in size, they were enriched in LCRs. For these analyses, we used as an index the LCR length in amino acids, versus the corresponding total protein length (**Fig. 7A**; **Supplemental File 7** for proteins with enriched LCRs). This result is consistent with the increase of RNA–binding proteins in PBs during HS (**Supplemental File 1**), which are known to contain LCRs in their RNA– interacting interfaces (Zagrovic et al., 2018). Hence, although disordered has been suggested as key for LLPS we did not find that in mock conditions PBs are richer in disorder than the corresponding controls.

**Figure 7.**
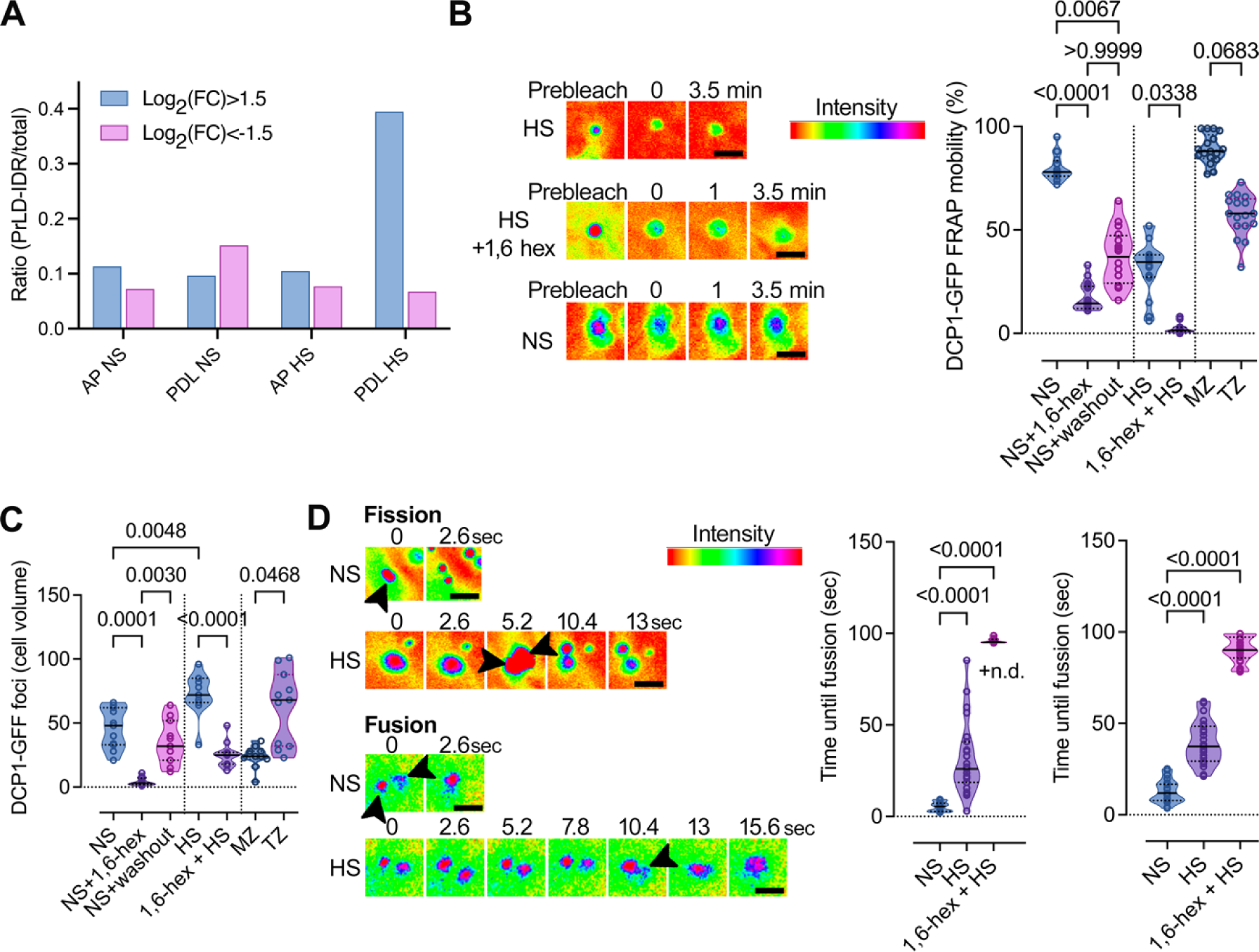
Content of PBs in proteins with LCRs during NS or HS and regulation of their dynamics. **A.** PrLDs and IDRs (LCRs) composition of APEAL (proteome)-enriched (log_2_FC>1.5)/excluded(log_2_FC<-1.5) proteins [from (Liu et al., 2023)]. The ratio here represents the LCR length sum (PrLDs+IDRs) divided by the total protein length. The PDL step can identify the disordered part of the proteome in PBs (Liu et al., 2023). AP, affinity purification; PDL, proximity-dependent ligation. **B.** FRAP assays in the presence or absence of 1,6-hexanediol (“hex”) in NS or HS from wild type (WT) expressing *DCP1pro:DCP1-GFP*. The experiment was replicated more than ten times. Note the lack of FRAP in the 1,6-hexanediol-treated sample. Right: relevant quantifications of mobile DCP1-GFP fraction from FRAP in the presence or absence of 1,6-hexanediol in NS or HS (and in washout experiments), and in MZ (meristematic) and TZ (transition) zones of the root. Data are means±SD (N=6 biological replicates with n=3 randomly selected meristematic epidermal cells 5 DAG each), and significance was calculated by an unpaired t-test (versus the NS; two-tailed *P* values are indicated). Scale bars, 400 nm. **C.** Quantifications of PBs in the presence or absence of 1,6-hexanediol (“hex”) in NS or HS from WT expressing *DCP1pro:DCP1-GFP* (cell volume in μm^3^). Data are means±SD (N=3 biological replicates with n=20 randomly selected meristematic epidermal cells 5 DAG each); significance determined by unpaired t-test (versus the NS; two-tailed *P* values are indicated). **D.** Super-resolution spinning disc microscopy images (combined with image deconvolution, ∼120 nm axial resolution at maximum acquisition speed of 0.1 sec per frame) of DCP1 fusion and fission dynamics, in NS or HS conditions from WT expressing *DCP1pro:DCP1-GFP*. Arrowheads indicate PBs; in fission, the two produced PBs are indicated for HS, while in fusion the arrowhead indicates in HS the coalescence. Scale bars, 200 nm. Right: quantification of corresponding fusion and fission events in the presence or absence of 1,6-hexanediol (“hex”) in NS or HS (N=10 biological replicates with n=3 randomly selected meristematic epidermal cells 5 DAG each; for 1,6-hexanediol N=2 biological replicates with n=1 randomly selected meristematic epidermal cells 5 DAG each; significance determined by Mann-Whitney). n.d., non-detected.

As HS increased the LCR content of PBs, we assumed that this could lead to a change in their properties. We compared the properties of large versus small PBs during NS or HS and in different root epidermal zones (i.e., meristematic vs. differentiating) through FRAP and upon 1,6–hexanediol treatments. Hence, PBs under HS (or large ones, >300 nm) showed reduced FRAP rates and partial insensitivity to 1,6–hexanediol (**Fig. 7B, C**). To further confirm this finding, we followed the fusion/fission rate of PBs in both NS and HS, as high rates of these two parameters indicate high liquidity (Linsenmeier et al., 2022). Accordingly, cell differentiation (i.e., transition zone) or HS significantly increased the time needed for fusion and fission suggesting that HS (and thus size) associates with reduced liquidity of PBs (**Fig. 7D**). Furthermore, as HS induced an increase of proteins in PBs containing canonical RNA–binding domains (**Supplemental File 1**), a combination of increased RNA–binding capacity, RNA content, and LCRs could contribute to PBs solidification. We propose that the IDR content of PBs associates well with their stability and RNA storability. This suggestion is also consistent with the increased disorder content at RNA–binding interfaces.

### RNAs can localize together with their encoded proteins in PBs

During the analysis of LCR content and its link to RNA storability, we serendipitously observed an enrichment of the m^6^A–readers YTH–homologs *EVOLUTIONARY CONSERVED C–TERMINAL REGION* (*ECTs*) mRNAs in PBs and at the same time, the corresponding homologous encoded proteins [**Supplemental File 8** (Liu et al., 2023)]. Intriguingly, this observation is reminiscent of observations in animals where many RNAs and their corresponding encoded proteins are stored in PBs (Hubstenberger et al., 2017). Hence, our observation prompted us to examine whether plant PBs have also the ability to store RNAs/cognate proteins and to what extent. As was done previously (Hubstenberger et al., 2017), we integrated the datasets for PBs proteome from APEAL (PBs–enriched proteins) and T–RIP (PBs–enriched RNAs) using the GOs “biological processes” or “cellular components”. We next ran a GO analysis of the common proteins/cognate RNAs (**Fig. 8A, B**). We evaluated hierarchical linkages between these terms using STRING (Search Tool for the Retrieval of Interacting Genes/Proteins; FDR < 0.05). Through GO–STRING we assigned hubs of cell wall remodeling and membrane remodeling, and wounding/ethylene responses associated with secondary metabolism for defense; as expected, we identified a high enrichment of PBs–related GOs (more than the ones observed for subnetworks above) related to RNA/nucleic acid metabolism and decapping [**Fig. 8A, B** (green)**, Supplemental Fig. 12A, B**]. On the other hand, we could not identify a similar enrichment of proteins/cognate RNAs from APEAL and T–RIP datasets using depleted proteins/RNAs or when using a permutated analysis between enriched/depleted datasets or with randomly enriched RNA/protein molecules. Examples of highly enriched cognate RNA/proteins related to actin–dynamics [e.g., Actin Depolymerizing Factors (ADFs), log_2_FC =3.95; see also *ADF9* above], veins and associated peptides [e.g., CLAVATA3/Embryo Surrounding Region–Related (CLE), log_2_FC=2.7], and cell wall [e.g., pectin methyl–esterase inhibitor 9 (PMEI9), log_2_FC=2.5 (Sorek et al., 2015)]. We also noticed GOs related to pectin, diverse (a)biotic stresses (i.e., bacterial, fungi, insects, heat, and salt stress) and a wounding/regeneration cluster with processes related to immunity and ethylene. These GOs are consistent with the phenotypes reported so far for plant decapping mutants, relevant to actin dynamics (e.g., trichome defects), veins, and cell wall remodeling (Xu et al., 2006; Xu and Chua, 2009).

**Figure 8.**
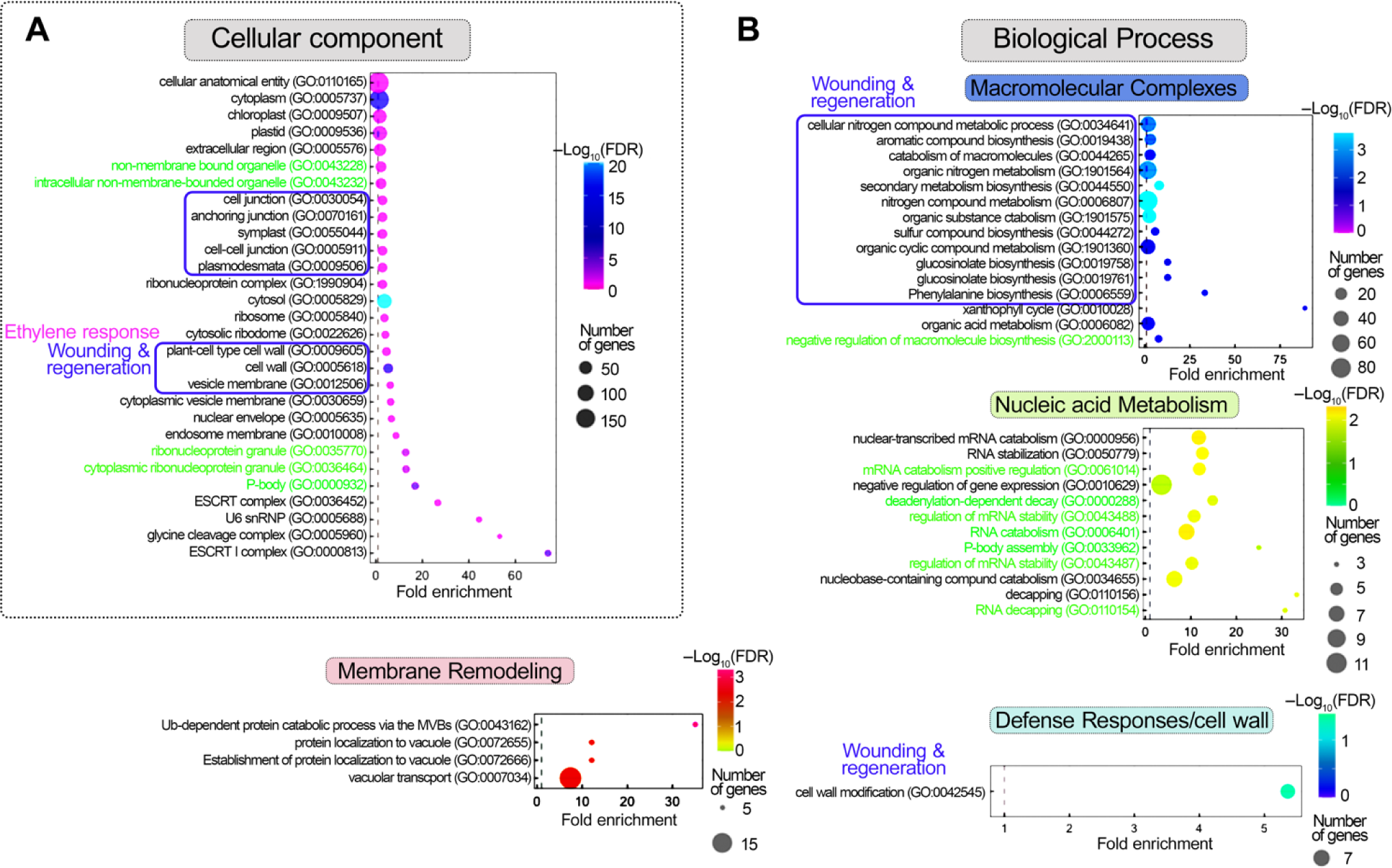
Gene ontologies identified for common proteins/cognate RNAs in PBs. **A.** and **B.** Cellular component (A) and Biological Process GO analysis (B), of common proteins/cognate of PBs-enriched RNAs and proteins. Co-cluster analyses of APEAL and T-RIP enrichments reveal hubs of cell wall remodeling, membrane remodeling, and wounding/ethylene responses associated with secondary metabolism for defense and RNA metabolism (GOs denoted with green color). FDR, false discovery rate.

We next queried whether cognate RNA/protein interactions in PBs depend on the affinity of certain amino acids to codons enriched in pyrimidines (PYR) or purines (PUR). The direct interactions between codons (in RNA) and the amino acids they code for have been proposed as a potential mechanism that drives cognate RNA/protein interactions (Zagrovic et al., 2018). These interactions are based on a proposed affinity scale based on PUR/PYR–amino acids interactions. Moreover, the lack of protein order (an increase of LCRs) during HS motivated further this analysis; residues in LCRs are not occupied to hold a rigid 3D structure but could be free for interactions with codons (Zagrovic et al., 2018). Yet, we did not observe differences between PYR/PUR matching with distributions of affine residues in IDRs or other protein segments when comparing PBs–enriched/non–enriched RNAs/protein pairs (**Supplemental Fig. 13A– C**). As a cautionary note though, affinities of residues and codons may become more significant in the context of crowded and viscous (more solid), dehydrated, low– dielectric environments like those present in PBs. Furthermore, LCRs in RNA–binding proteins may promote the association of RNAs with PBs. In accordance, we observed that RNA targets of the RBP rich in LCRs GRP7 [Gly Rich Protein 7, AT2G21660, (Meyer et al., 2017)] that localizes in PBs (Liu et al., 2023), could also be found in PBs (**Supplemental File 10**). Our results thus suggest that the recruitment of RNA–binding proteins and the increase of LCRs in PBs associate with a marked increase in RNAs. The exact mechanism by which RNAs and cognate proteins localize in PBs, while others escape from PBs (in HS), merits further investigation.

### The SCAR/WAVE–DCP1 axis can regulate RNA dynamics

In non–plants, PBs compete with polysomes in a complex manner for RNAs (Cougot et al., 2013). Accordingly, we observed that most RNAs stored in PBs encode proteins and thus could in principle affect translation (**Fig. 9A**; both in NS and HS, 95% in PBs versus 80% in the total transcriptome). As PBs store mainly mRNAs, we aimed to evaluate whether PBs could affect their translation dynamics. As we have previously shown that the dissolution of PBs is at least partially controlled by the SCAR/WAVE– DCP1 axis, we assumed that this axis could modulate the PBs–to–translation shuttling for at least a portion of PBs–enriched RNAs (**Fig. 9B**, model). We thus queried whether PBs–enriched RNAs from seedlings of wild type, the *scar1234* (more/larger PBs) or the weak allele *dcp1–3* (fewer PBs), are depleted from polysomes where RNAs are translationally active (Jang et al., 2019). To isolate polysomes, we refined a ribosome profiling approach on a sucrose cushion with 12 fractions (Yanguez et al., 2013). Density ribosome profiles of *scar1234*, and *dcp1–3* matched that of the wild type (**Supplemental Fig. 14A, B**; A_260_), suggesting that profiling would be rather specific and not reflect a broad translational regulation, as observed for other mutants (Cho et al., 2022). We used RT–qPCR in combined fractions of monosomes (first 4 fractions of the 12 collected in total) and polysomes (fractions 5–12) to determine levels of mRNAs in wild–type, *scar1234*, and *dcp1–3,* comparing them to the total mRNA of the same samples corresponding to the 10% (v/v) of the initial polysome profiling extract (**Fig. 9C, D**). We selected *EBF2*, *DFL1* [*DWARF IN LIGHT 1, GH3.6, GRETCHEN HAGEN3.6*)*, RAP2.4, RAP2.4D, AAK6 (ARABIDOPSIS ADENYLATE KINASE 6)*, and *PMEI9*; **Supplemental File 11**], and for normalization, we used *TUBULIN ALPHA–4 CHAIN* (AT1G04820) that was excluded from PBs and showing similar equivalent levels in all fractions (**Supplemental Fig. 14C**). We observed a relatively good correlation between polysome occupancy and PBs–depletion for some RNAs but not for all (**Fig. 9D**, e.g., AAK6).

**Figure 9.**
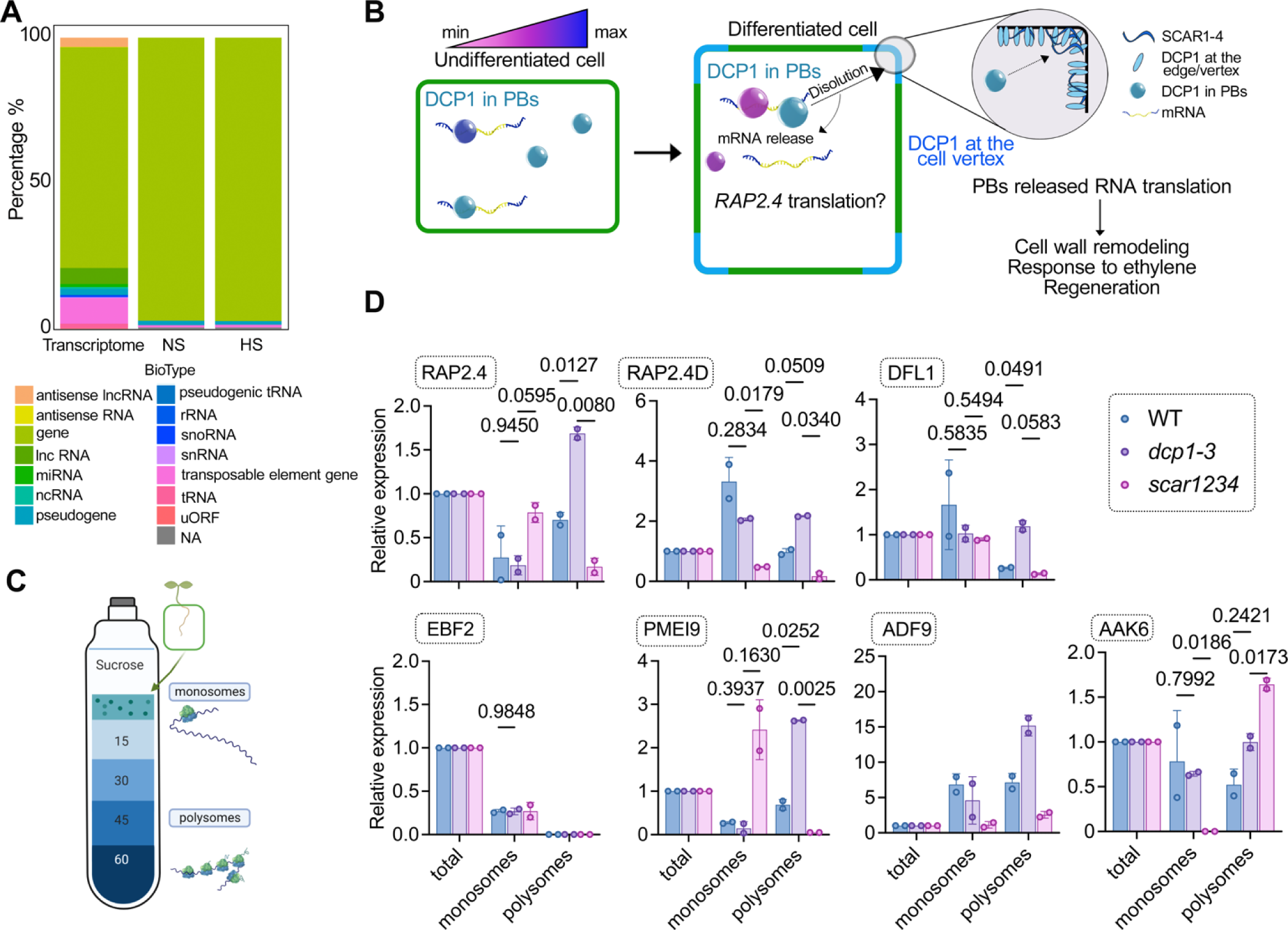
Regulation of translational dynamics by the SCAR/WAVE–DCP1 axis. **A.** RNA classes of PBs-enriched RNAs (%). Total RNA corresponds to the distribution of RNAs in the whole transcriptome. NA, non-available or not defined. Lnc, long non-coding; mi, micro; nc, non-coding; sno, small nucleolar; uORF, upstream open reading frame. **B.** Model for the dissolution of PBs by the SCAR/WAVE–DCP1 axis. The SCAR/WAVE (mainly through SCAR2) entraps DCP1 at the edge/vertex (through SCAR2), which leads to PBs dissolution and consecutive mRNA release for translation, e.g., of *RAP2.4*. Processes like cell wall remodeling-related processes, such as response to ethylene and regeneration could be affected. The colour-coding for PBs corresponds to their ability for RNA decay. **C.** Cartoon showing the polysome profiling approach. The denser part [60% (w/v) sucrose] corresponds to RNAs associated with polysomes where they are mostly translated (Jang et al., 2019). **D.** RT–qPCR of *RAP2.4, APUM, EBF2*, *DFL1*, and *PMEI9* from total RNA, monosome, or polysome fractions in wild type (WT), *dcp1-3*, or *scar1234*. Data are means of two independent experiments with two technical replicates (n=2 RT–qPCR assays). The data were normalized against *TUBULIN* and the levels of RNAs in the input which is denoted as “total”; significance was determined by two-way ANOVA using a Geisser-Greenhouse correction (due to unequal variance) and Fisher’s exact test for multiple comparisons.

**Figure 10.**
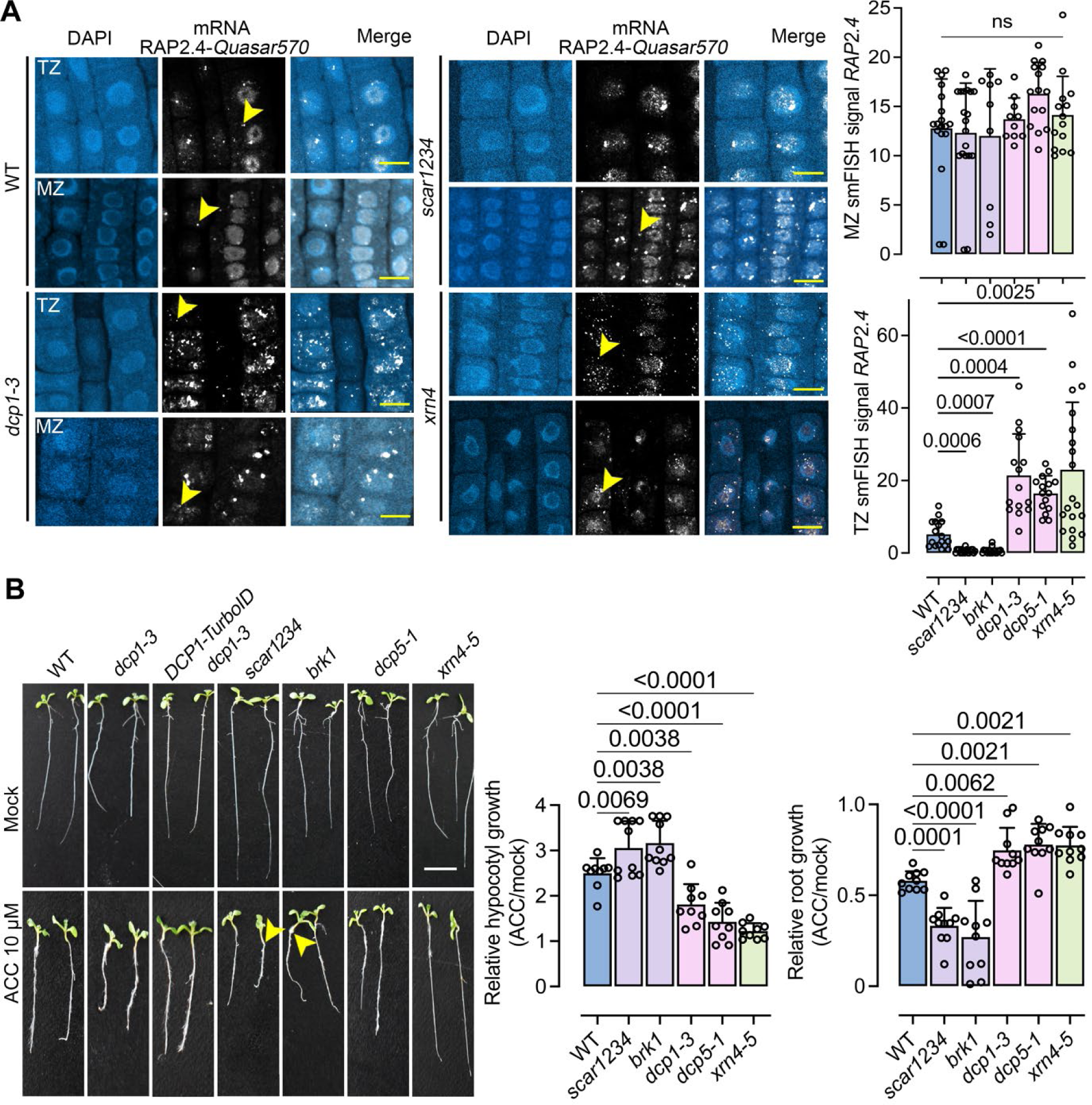
Modulation of RAP2.4 levels and ethylene response by the SCAR/WAVE–DCP1 axis. **A.** Micrographs from smFISH for detection of *RAP2.4*. mRNA (grey) in the root epidermal cells of the meristematic (MZ) and transition (TZ) zones in the corresponding mutants [wild type (WT), *dcp1-3, scar1234,* and *xrn4-5*]. Insets: DCP1–GFP/*RAP2.4* puncta colocalization. The yellow arrowheads denote *RAP2.4* signal. The experiment was replicated three times. Scale bars, 12 μm. Right: corresponding quantifications with the number of smFISH spots in WT, *dcp1-3, scar1234,* and *xrn4-5*. Data are means of three independent experiments with two technical replicates each (n=2 roots each with 10 MZ or TZ randomly selected cells 5 DAG), and significance was determined by ordinary one–way ANOVA. ns, non-significant. **B.** Photos showing growth in the absence or presence of the ACC ethylene precursor (10 μM 5 DAG). The experiment was replicated three times (n=10 seedlings in each replicate). Arrowheads denote the elongated hypocotyls in *scar1234* and *brk1*. Also, note the complementation of DCP1-TurboID expressing *dcp1-3* (mainly upon ACC treatment). Scale bars, 1 cm. Right: corresponding quantifications of hypocotyl and root length in the presence or absence of ACC. Data are means ±SD (N=3 biological replicates with n=10 roots/hypocotyls each); significance was determined by ordinary one–way ANOVA with Dunnett’s corrections.

In this context, we decided to further study *RAP2.4* as it is rapidly decayed through the decapping complex [results herein and (Sorenson et al., 2018)], showed enrichment in GFP–trap experiment, correlated well with PBs–depletion/translation, and fits in the central PBs subnetwork and RNA/cognate protein hub identified, “response to wounding/regeneration” and “ethylene” [**Fig. 2** for GO analysis and **Fig. 7**; (Lin et al., 2008; Iwase et al., 2011)]. According to the polysome and the decay results, smFISH signal accumulation *RAP2.4* inversely correlated to that of the SCAR/WAVE–DCP1– axis which is more active at the root transition zone onwards (i.e., dissolving PBs; **Fig. 10A**). In more detail, *dcp1–3* (or *dcp5–1*) had increased cytoplasmic levels of *RAP2.4* mRNA throughout the root, while *scar1234* had reduced *RAP2.4* mRNA levels at the transition zone compared to the wild type. We though observed high nuclear levels of *RAP2.4* in *scar1234*, suggesting that high decay rates of *RAP2.4* feedback to transcription, as suggested for other RNAs in plants (Manavella et al., 2023). *RAP2.4* mRNA decay depended on XRN4, as in the loss–of–function *xrn4–5* mutants *RAP2.4* mRNA was increased (**Fig. 10A**). Overall, these results suggest that an RNA–specific combination of storage and decay by PBs modulate some RNA levels channeled into translation. As PBs can exchange RNAs with SGs, and as in non–plants it has been shown that translation can take place, we assume that storage of RNAs in PBs could also enhance translation independently of polysomes (Pitchiaya et al., 2019; Baumann, 2021). Furthermore, decay rates can feedback to transcription and translation, a finding that merits further investigation as to which are the underlying mechanisms.

### The SCAR/WAVE–DCP1 axis modulates ethylene signaling

Given the link of *RAP2.4* with ethylene, we asked whether the SCAR/WAVE–DCP1– axis modulates ethylene signaling. PBs sequester and degrade *EBF1* and *EBF2* through XRN4 [also known as ETHYLENE INSENSITIVE 5; (Li et al., 2015; Merchante et al., 2015)]. Accordingly, *EBF2* RNA in *dcp1–3* was increased as has also been reported for *xrn4* (by 50%, as both mutants show reduced RNA decay). As predicted above using STRING–GO, we observed that ethylene or HS led to the localization in PBs of both the cognate RNA/protein *RAP2.4* pair (**Supplemental Fig. 15A, B**). Furthermore, ethylene could induce the removal of SCAR/WAVE from the vertex in the transition zone (and above) of the root leading to an expected increase in the size/number of PBs (**Supplemental Fig. 16**). We speculate for now that this removal depends on the ethylene–induced activation of constitutive photomorphogenic 1 (COP1) E3 ligase, which degrades the SCAR/WAVE (Dyachok et al., 2011). Accordingly, *scar1234* and *brk1* (*BRICK1*), another SCAR/WAVE component mutant, were sensitive to applications of ethylene precursors ACC or Ethephon [(2–Chloroethyl)phosphonic acid, known also as “Ethrel”] determined as increased hypocotyl elongation under light conditions [as the *rap2.4* mutant, (Lin et al., 2008)], unlike decapping mutants showing reduced sensitivity (**Fig. 10B, C**). These results provide a long–sought link between ethylene modulation of PBs number and activity and extend our understanding of ethylene signaling.

Collectively, we present approaches for the study of RNA composition, properties, and activities of condensates in plants. We further suggest that certain properties of PBs regulate their ability to store RNA, thereby remodeling the proteome through the translation and decay of selected RNAs. In animals, the fragile X mental retardation protein (FMRP, neuron generation) inhibits translation by binding to accessory proteins of the SCAR/WAVE (Kim et al., 2019). Another FMRP member form condensates sequestering RNA and the translational machinery to activate translation (Kang et al., 2022). We thus offer a different model for translational regulation by the SCAR/WAVE.

## METHODS

### Plant material

All the plant lines used in this study were in the Arabidopsis Columbia–0 (Col–0) ecotype or *Nicotiana benthamiana* unless stated. Primers used for genotyping and cloning are described in **Supplemental Table 1**. The following mutants and transgenic lines used in this study were described previously: *dcp1–1* (Xu et al., 2006), *dcp1–3* (Martinez de Alba et al., 2015), *dcp2–1* (Chantarachot et al., 2020), *dcp5–1* (Xu and Chua, 2009), *xrn4–5* (Souret et al., 2004), *brk1* (Dyachok et al., 2008), *scar1234* (Dyachok et al., 2008), *35Spro:GFP–DCP1* (Gutierrez-Beltran et al., 2015), *35Spro:DCP2–YFP* (Jang et al., 2019)*, fip37–4* (Bodi et al., 2012) and *DCP1pro:DCP1– GFP*, *SCAR2pro:mCherry–SCAR2* (Chin et al., 2021). Seedlings carrying the TurboID (*35Spro:sGFP–TurboID–HF*) construct have been previously described (Liu et al., 2023). In all experiments, plants from T1/F1 (co–localization experiments), T2/F2, or T3 (for physiological experiments) generations were used.

### Plant growth, and pharmacological treatments

Arabidopsis seedlings were sterilized and germinated on half–strength Murashige and Skoog (MS) agar medium under long–day conditions (16 h light/8 h dark). In all experiments involving the use of mutants or pharmacological treatments, the medium was supplemented with 1% (w/v) sucrose or as otherwise specified. Arabidopsis plants for crosses, phenotyping of the above–ground part, and seed collection were grown on soil (comprising of a blend of sphagnum moss peat, compost, worm castings, perlite, and a special ‘natural plant booster’) in a plant chamber at 22°C/19°C, 14–h–light/10–h– dark or 16–h–light/8–h–dark cycles, and light intensity 150 µE m^−2^ s^−1^. *Nicotiana benthamiana* plants were grown in Aralab or Percival cabinets at 22°C, 16–h–light/8–h– dark cycles, and a light intensity of 150 µE m^−2^ s^−1^. Vertically grown 4 to 5–d–old Arabidopsis seedlings were transferred and incubated in a half–strength liquid MS medium containing corresponding drugs for each specific time course treatment as indicated. Ethephon treatment was performed in a final concentration of 100 μM (from a 100mM stock) in seedlings or *N. benthamiana* slices for 20 minutes. Long ethephon or 1–aminocyclopropane–1–carboxylate (ACC) treatment was performed on half–strength MS plates in a 10 μM final concentration (under light or in the dark). For 1,6–hexanediol treatments, 10 % (v/v) aqueous solution was used. The stock solutions of 50 mM biotin, 100 mg/ml cycloheximide (CHX), were dissolved in DMSO. Cordycepin treatments were performed in a 1 mM final concentration in incubation buffer (15 mM sucrose, 1 mM piperazine–N, N′–bis(2–ethane sulfonic acid (PIPES) pH 6.25, 1 mM KCl, 1 mM sodium citrate) (Sorenson et al., 2018).

### Bacterial strains, clonings, and constructs

Cloning of RAP2.4pro:RAP2.4–venusYFP was done according to standard In–fusion cloning procedures (Takara In–Fusion® Snap Assembly Master Mix #638948), using the linearized pGWB601 vector. Primers used for this cloning can be found in **Supplemental Table 1.** The E. coli strains NEB10 (New England Biolabs #C3019H) or NEB stable (New England Biolabs #C3040H; CRISPR constructs) were used for clonings. Electrocompetent *Agrobacterium tumefaciens* C58C1 Rif^R^ (pMP90) or GV3101 Rif^R^ bacterial cells (i.e., a cured nopaline strain commonly used for tobacco infiltration) were used for electroporation and tobacco infiltration.

### Transient transformation of Nicotiana benthamiana

1. *N. benthamiana* plants were grown under normal light and dark regimes at 25°C and 70% relative humidity. Three– to four–week–old *N. benthamiana* plants were watered from the bottom ∼24 h before infiltration. Transformed Agrobacterium strain C58C1 Rif^R^ (pMP90) or GV3101 Rif^R^ harboring the constructs of interest were used to infiltrate *N. benthamiana* leaves and for transient expression of binary constructs by Agrobacterium–mediated transient infiltration of lower epidermal leaf cells. Transformed Agrobacterium colonies were grown for ∼20 h in a shaking incubator (200 rpm) at 28°C in 5 mL of yeast extract peptone (YEP) medium (10 g/L yeast extract, 10 g/L peptone, 5 g/L NaCl and 15 g/L bacterial agar), supplemented with appropriate antibiotics (i.e., 100 g/L spectinomycin). After incubation, the bacterial culture was transferred to 15–mL Falcon tubes and centrifuged (10 min, 3,500 g, RT). The pellets were washed with 5 mL of infiltration buffer (10 mM MgCl_2_, 10 mM MES pH 5.7), and the final pellet was resuspended in infiltration buffer supplemented with 200 μM acetosyringone. The bacterial suspension was diluted with infiltration buffer to adjust the inoculum cell density to a final OD_600_ value of 0.4. The inoculum was incubated for 2 h agitating at 28°C before infiltration into *N. benthamiana* leaves by gentle pressure infiltration of the lower epidermis of leaves (fourth and older true leaves were used, and about 4/5–1/1 of their full size) with a 1–mL hypodermic syringe without a needle.

### T–RIP method

All materials were RNase–free. Seven–day–old seedlings were submerged in 50 ml of 50 μM biotin (in 10 mM MgCl_2_, 10 mM MES pH 5.7) for 1 min under vacuum. After a 24 h incubation on a new ½ MS plate, the plates were treated with heat stress (37 °C, 2 h). Around 1–2 g of seedlings’ fresh weight was harvested in 1% (v/v) formaldehyde (FA) solution and a 15 min vacuum was applied. FA was quenched by adding glycine to a final concentration of 125 mM for 5 min under vacuum. The plant material was rinsed 2× with RNase–free H_2_O and pulverized in liquid N_2_. Proteins were extracted as described for APEAL (Liu et al., 2022); the extraction buffer was supplemented though with a 20 U/ml RNase inhibitor (ThermoFisher Scientific, EO0381). The supernatant was loaded on a PD–10 column (Cytiva, GE17–0851–01) for the removal of the excess biotin and incubated with equilibrated 100 μL Dynabeads™ MyOne™ Streptavidin (ThermoFisher Scientific, 65601) at 4 °C for 2 h with gentle rotation. The beads were precipitated using a magnetic rack and washed four times with extraction buffer and two times with dilution buffer (extraction buffer without detergent). Then RNA–protein complexes were eluted from the beads with 100 μL protease buffer (0,5 mg/ml Proteinase K, 20 U/ml RNase inhibitor in 30 mM Tris–HCl, pH 8.0) at 55 °C for 30 min. RNA was extracted from the eluted beads with phenol: chloroform: iso–amyl alcohol (25:24:1) (v/v/v) (VWR, 136112– 00–0), followed by precipitation with 1:1 (v/v) isopropanol with 0.1 volume of 3M NaAc, pH 5.2 and 2 μL Glycoblue (ThermoFisher Scientific, AM9515) for 1 h. After centrifugation at 4 °C, 16,000 g for 20 min, the blue, due to Glycoblue, pellet was washed 2× with 75% (v/v) ethanol. Air–dried pellet was resuspended into 15 μL of DNase–, RNase–, and protease–free water. The concentration of RNA was determined by Qubit^®^ RNA HS Assay Kit (New England BioNordika BioLab, Q32852). All the RIP– RNA samples were treated with DNase I (ThermoFisher Scientific, EN0521) and Ribosomal RNA depletion (RiboMinus™ Plant Kit (ThermoFisher Scientific, A1083808).

The RNA was measured with Qubit® RNA HS Assay Kits again and libraries were prepared with NEBNext^®^ Ultra™ II RNA Library Prep with Sample Purification Beads (Invitrogen Life Technologies (Ambion Applied Biosystem), E7775S) and NEBNext^®^ Multiplex Oligos for Illumina^®^ (Dual Index Primers Set 1) (New England BioNordika BioLab, E7600S). cDNA library quality was monitored with Agilent DNA 7500 Kit (Agilent Technologies Sweden AB, 5067–1506). cDNA libraries were sequenced with a paired–end sequencing strategy to produce 2 × 150–bp reads using Novogen sequencers (Novogene, England).

### GFP–RIP method

A modified T–RIP protocol was followed. Following fixation and formaldehyde quenching steps, as described for T–RIP, the lysates of 7–day–old seedlings of *35Spro:DCP1–GFP, DCP1pro:DCP1–GFP/dcp1–1, 35Spro*:TAP–GFP (NS and HS) was added on ChromoTek GFP–Trap® Agarose beads (Cat No. gta) for 2 h with gentle rotation at 4 °C. Beads were precipitated and washed with centrifugation at 1,000g for 2 min. After the Proteinase K treatment and RNA extraction, RNA samples were cleaned up with the Monarch® RNA Cleanup Kit (50 μg, T2040L). Reverse–transcriptase reaction was performed using the Minotech RT (Catalogue No. 801–1(10KU)) kit and the qPCR using the SYBR™ Fast Green Master mix (Thermo Fisher, 4385612).

### Immunoblotting

Infiltrated *N. benthamiana* tobacco leaves or Arabidopsis leaves and seedlings were harvested, and proteins were extracted. The tissue samples were flash–frozen in liquid Ν_2_ and kept at –80°C until further processing (not more than a month). The samples were crushed using a liquid Ν_2_–cooled mortar and pestle and the pulverized powder was transferred to a 1.5 ml tube. Extraction buffer (EB; 50 mM Tris–HCl (pH 7.5), 150 mM NaCl, 10% (v/v) glycerol, 2 mM Ethylene diamine tetraacetic acid (EDTA), 5 mM Dithiothreitol (DTT), 1 mM phenylmethylsulphonyl fluoride (PMSF), Protease Inhibitor Cocktail (Sigma–Aldrich, P9599), and 0.5 % (v/v) octyl phenyl–polyethylene glycol (IGEPAL CA–630, Sigma–Aldrich) was added. The lysates were pre–cleared by centrifugation at 16,000 g at 4 °C, for 15 min, and the supernatant was transferred to a new 1.5 ml tube. This step was repeated 2× and the protein concentration was determined by the RC DC Protein Assay Kit II (Bio–Rad, 5000122). Laemmli buffer was added at a 1:2 ratio, and equivalent amounts of protein (∼30 μg) were separated by sodium dodecyl sulfate–polyacrylamide gel electrophoresis (SDS–PAGE) (1.0 mm thick 4 to 12% gradient polyacrylamide Criterion Bio–Rad) in 3–(N–morpholino) propane sulfonic acid (MOPS) buffer (Bio–Rad) at 150 V. Subsequently, proteins were transferred onto a Polyvinylidene fluoride (PVDF; Bio–Rad) membrane with 0.22 μm pore size. The membrane was blocked with 3% (w/v) BSA fraction V (ThermoScientific) in Phosphate Buffer Saline–Tween 20 (PBS–T) for 1 h at room temperature (RT) followed by incubation with horseradish peroxidase (HRP)–conjugated primary antibody at RT for 2 h (or primary antibody for at RT 2 h followed by the corresponding secondary antibody at RT for 2 h). The following antibodies were used: streptavidin– HRP (Sigma–Aldrich; 1:25,000), and FLAG–HRP (Sigma–Aldrich, A8592, 1:2,000). Chemiluminescence was detected with ECL Prime Western Blotting Detection Reagent (Cytiva, GERPN2232) and SuperSignal™ West Femto Maximum Sensitivity Substrate (ThermoFisher Scientific, 34094).

### Detection of RNA molecules by whole–mount–smFISH

The mRNA probes used were selected following the criteria 1) roughly 18–22 bases long, and 2) GC content close to 45%. Furthermore, individual probes within a set had at least two nucleotides of space between their target regions. The Stellaris Probe Designer version 4.2 (Biosearch Technologies) was used for probe design and ordering (https://www.biosearchtech.com/stellaris-designer). Localization was performed following the protocols from Stellaris RNA fish (Protocol for Arabidopsis root meristem) and (Duncan et al., 2016). The roots of 4 to 5–day–old Arabidopsis seedlings were fixed for 2 h in fixation buffer (50 mM piperazine–N,N′–bis(2–ethane sulfonic acid (PIPES), pH 7.2, with 20 mM EGTA (ethylene glycol–bis(β–aminoethyl ether)–N,N,N′, N′– tetraacetic acid) and 20 mM MgSO_4_ containing 2% w/v paraformaldehyde, 0.1% Triton X–100 and 400 μM Maleimidobenzoyl–N–hydroxy succinimide ester (ThermoFisher Scientific, 22311), washed two times with 1x Phosphate–buffered saline (PBS) buffer and then incubates four–days with ClearSee solution (xylitol powder [10% (w/v)], sodium deoxycholate [15% (w/v)] and urea [final 25% (w/v)] in Rnase–free water) (Kurihara et al., 2015). Then the seedlings were washed 2× with 1× Phosphate–buffered saline (PBS) buffer and followed with wall digestion at 37 °C for 30 min [in 0.2% (w/v) driselase and 0.15% (w/v) macerozyme]. The roots were washed two times with washing buffer (50 mM piperazine–N,N′–bis(2–ethane sulfonic acid (PIPES), pH 7.2, with 20 mM EGTA (ethylene glycol–bis(β–aminoethyl ether)–N,N,N′,N′–tetraacetic acid) and 20 mM MgSO_4_) and incubated in permeabilization buffer (3% v/v octylphenyl– polyethylene glycol IGEPAL CA–630,10% dimethyl sulfoxide, 50 mM piperazine–N,N′– bis(2–ethane sulfonic acid (PIPES), pH 7.2, with 20 mM EGTA and 20 mM MgSO_4_) at 37 °C for 30 min. Then set samples were washed 2× with wash buffer (10% formamide and 2 × saline–sodium citrate (SSC)), 5 min per wash. Then slides were incubated with 100 μL 250 nM of probes (in hybridization solution: 10 % dextran sulfate, 2 × saline– sodium citrate (SSC), and 10% formamide) at 37 °C for 4 h or overnight hybridization at 4 °C in the dark. Samples were washed 2× with wash buffer for 5 min each. Slides were incubated with 100 μL nuclear stain 4,6–diamidino–2–phenylindole (DAPI; 100 ng ml^−1^) for 30 min at 37 °C in the dark. Excess DAPI was removed and 100 μL 2 × saline– sodium citrate (SSC) was added and then washed away. A drop of Vectashield antifade mounting medium was added and sealed with coverslip and nail polish. A Zeiss 800 confocal microscope was used for imaging, using a 63x NA=1.4 water–corrected immersion objective, and the wide–field mode was used to obtain all images in standard settings. The following wavelengths were used for fluorescence detection: for probes labeled with Quasar 570 an excitation line of 561 nm was used and the signal was detected at wavelengths 570–640 nm; for DCP1–GFP an excitation line of 488 nm was used and the signal was detected at wavelengths of 530–550 nm; for DAPI an excitation line of 405 nm was used and the signal was detected at wavelengths of 420–460 nm.

### Quantification of fluorescent intensity, FRAP, and FRET

To create the most comparable lines to measure the fluorescence intensity of reporters in multiple mutant backgrounds, we crossed homozygous mutant bearing the marker with either a WT plant (outcross to yield progeny heterozygous for the recessive mutant alleles and the reporter) or crossed to a mutant only plant (backcross to yield progeny homozygous for the recessive mutant alleles and heterozygous for the reporter).

Fluorescence was measured as a mean integrated density in regions of interest (ROIs) with the subtraction of the background (a proximal region that was unbleached and had less signal intensity than the signal of the ROI region). FRAP mode of Zeiss 780 ZEN software was set up for the acquisition of 3 pre–bleach images, one bleach scan, and 96 post–bleach scans (or more). Bleaching was performed using a 488 nm laser line at 100% transmittance and 20–40 iterations depending on the region and the axial resolution (iterations increased in deeper tissues to compensate for the increased light scattering). In FRAP the width of the bleached ROI was set at 2–10 µm. Pre– and post– bleach scans were at minimum possible laser power (0.8 % transmittance) for the 458 nm or 514 nm (4.7%) and 5% for 561 nm; 512 x 512 8–bit pixel format; pinhole of 181 μm (>2 Airy units) and zoom factor of 2.0. The background values were subtracted from the fluorescence recovery values, and the resulting values were normalized by the first post–bleach time point and divided by the maximum fluorescent time–point set maximum intensity as 1. The objective used was a plan–apochromat 20x with NA=0.8 M27 (Zeiss). The following settings were used for photobleaching DCP1: 10–20 iterations for DCP1–GFP; 10 to 60 s per frame; 100% transmittance with the 458– to 514–nm laser lines of an argon laser. Pre– and post–bleach scans were at the minimum possible laser power (1.4 to 20% transmittance) for the 488 nm and 0% for all other laser lines, 512×512 pixel format, and zoom factor of 5.1. The fluorescence intensity recovery values were determined, then the background values were subtracted from the fluorescence recovery values, and the resulting values were normalized against the first post–bleach time point. FRET analyses were done using the method described previously (Liu et al., 2023).

### Proximity Ligation Assay (PLA) protocol

For PLA immunolocalization the primary antibody combinations diluted 1:200 for α– FLAG mouse (Sigma–Aldrich, F1804), and α–GFP rabbit (Millipore, AB10145). The antibodies were incubatedat 4°C. Roots were then washed with microtubule–stabilizing buffer (MTSB: 50 mM piperazine–N,N′–bis(2–ethane sulfonic acid (PIPES), 5 mM EGTA (ethylene glycol–bis(β–aminoethyl ether)–N,N,N′,N′–tetraacetic acid), 2 mM MgSO_4_, 0.1% [v/v] Triton X–100) and incubated at 37°C for 3 h either with α–mouse plus and α–rabbit minus for PLA assay (Sigma–Aldrich, 681 Duolink). PLA samples were then washed with MTSB and incubated for 3 h at 37°C with ligase solution as described (Pasternak et al, 2018). Roots were then washed 2× with buffer A (Sigma– Aldrich, Duolink) and treated at 37°C for 4 h in a polymerase solution containing fluorescent nucleotides as described (Sigma–Aldrich, Duolink). Samples were then washed 2× with buffer B (Sigma–Aldrich, Duolink), with 1% (v/v) buffer B for another 5 min, and then the specimens were mounted in Vectashield (Vector Laboratories) medium. Root tips were imaged using a Zeiss 780 confocal laser scanning microscope using a 63x NA=1.6 oil–corrected immersion objective.

### Ribosome profiling

A modified protocol described previously was used (Yanguez et al., 2013). All materials were RNase–free, and all buffers were autoclaved for 15 min. Two hundred mg of seedlings grown on half–strength MS plates were used as a starting material. Tissue lysis was performed with a modified Polysome Extraction Buffer (0.2 M Tris–Cl pH 9.0, 0.2 M KCl, 25 mM EGTA (ethylene glycol–bis(β–aminoethyl ether)–N,N,N′,N′– tetraacetic acid) 35 mM MgCl_2_, 1% detergent mix from a 20% stock solution; 20% (w/v) polyoxyethylene(23)lauryl ether (Brij–35), 20% (v/v) Triton X–100, 20% (v/v) octyl phenyl–polyethylene glycol (IGEPAL CA–630), 20% (v/v) polyoxyethylene sorbitan monolaurate 20 (Tween 20), 1% (v/v) Polyoxyethylene 10 tridecyl ether (PTE), 5mM DTT, 1mM PMSF, 50 μg/mL cycloheximide, 50 μg/mL chloramphenicol). The ultracentrifuge step was performed in 3ml final volume tubes and with a swing–out rotor sorvall TST60.4. RNA extraction was done with TriZol from the 12 fractions collected and then samples were cleaned up using the Monarch® RNA Cleanup Kit (50 μg) from New England Biolabs (NEB). Optical densities (OD_260_) were used to calculate RNA concentration. Reverse–transcriptase reaction was performed using the Minotech RT (Catalogue No. 801–1(10KU)) kit and the qPCR using the SYBR™ Fast Green Master mix (Thermo Fisher, 4385612).

### Decay levels determination

Five–day–old seedlings were treated with cordycepin (Sigma, C3394) according to (Sorenson et al., 2018) and for the following time points t_0_, 60 min and 240 min. RNA was extracted with the Spectrum™ Plant Total RNA Kit (Sigma, STRN250). Reverse– transcriptase reaction was performed using the Minotech RT (Catalogue No. 801– 1(10KU)) kit and the qPCR using the SYBR™ Fast Green Master mix (4385612).

### N^6^–methyladenosine (m^6^A) purification

The m^6^A immunoprecipitation was performed from 5–day–old seedlings using the EpiMark® N^6^–Methyladenosine Enrichment kit (NEB) starting from 4 μg of total RNA. Reverse–transcriptase was performed using the Minotech RT [Catalogue No. 801–1(10 KU)] kit and the qPCR using the SYBR™ Fast Green Master mix (Thermo Fisher, 4385612).

### Visualization of networks and analyses

For visualization, Cytoscape 3.5.1 was used. Tab–delimited files containing the input data were uploaded. The default layout was an edge–weighted spring embedded layout, with the NormSpec used as edge weight. Nodes were manually rearranged from this layout to increase visibility and highlight specific proximity interactions. The layout was exported as a PDF and eventually converted to a TIFF file.

### In silico Data analyses

For RNAs that are enriched in PBs, the log_2_ (DCP1/GFP) value (from the RNA–seq dataset) was used, as an indicator of the enrichment of each RNA with PBs. The “Fold_enrichment” values from the graphs, which are associated with m^6^A modifications, are referred to as the average “Fold_enrichment” value of each RNA from the given dataset (Parker et al., 2020). In brief, the *VIRILIZER* (*vir–1*) mutant is a conserved m^6^A component of the writer complex, and the m^6^A was compared to a line expressing VIR fused to Green Fluorescent Protein (GFP) (VIR complemented; *virC*) that restores VIR activity in the *vir–1* background. To provide the m^6^A sites, the differential error rate analysis approach was used (using the base–calling) to compare the mutant (defective m^6^A) and VIR–complemented lines. Through this approach, 17,491 sites with an FC of >2 error rate (compared with the TAIR10 reference base) in the VIR–complemented line were identified. The “Fold_enrichment” values indicate the statistical importance of the detected RNA to be methylated to one or more sites. Information about the total length of the GC% of 5’ and 3’ UTR has been retrieved from the TAIR database (https://www.arabidopsis.org/) and plotted against the corresponding log_2_(DCP1/GFP) values.

### Quantification and Statistics

All statistical data show the mean ± SD of at least three biologically independent experiments or samples, or as otherwise stated. *N* denotes biological replicates, and “*n*” technical replicates or population size. Statistical analyses were performed in GraphPad (https://graphpad.com/) or R studio (R-project.org). Each data set was tested whether it followed normal distribution when *N* ≥ 3 by using the Shapiro normality test. The significance threshold was set at *p*<0.05 (significance claim), and the exact values are shown in the graphs. Graphs were generated by GraphPad Prism or R. Details of the statistical tests applied, including the choice of the statistical method, are shown in the corresponding figure legend. In boxplots or violin plots, upper and lower box boundaries represent the first and third quantiles, respectively, horizontal lines mark the median and whiskers mark the highest and lowest values. Figures relating to sequence properties i.e., Codon Usage Bias, were created using custom Python (https://www.python.org/) and R scripts. Associated boxplots follow the same characteristics as stated previously and any overlying data points denote the included values. In histograms, vertical lines refer to the median of the presented distributions whereas boxplots, when present, are used as a visual alternative for statistical purposes. In scatterplots, existing lines represent the loess–smoothed representations of the underlying points. Kernel density plots were created using a Gaussian kernel. Finally, the height of related bar plots accounts for the median value in all cases. Displayed *P*–values in the case of multiple– comparison testing was calculated using a Dunn’s posthoc (Bonferroni; FDR<0.05) test after a statistically significant Kruskal–Wallis ANOVA (p–value<0.05). In the case of pairwise hypothesis testing a Mann–Whitney U test was used instead (p–value<0.05).

### Image analyses and preparation

Image analyses and intensity measurements (Integrated Density) were done using Fiji v. 1.52 software (rsb.info.nih.gov/ij). The intensities were normalized for relative values against the background signal intensity in raw images/micrographs. The calculations were done with arbitrary units (denoted as AUs). The dwell time rate of tagged proteins in FRAP experiments was calculated by the single exponential fit (Moschou et al., 2016; Deli et al., 2022). Colocalization was analyzed using Pearson Correlation Coefficient (PCC) statistics (Spearman or Manders analyses produced similar results (French et al., 2008). Images were prepared in Adobe Photoshop v. 2023. Time series movies were compressed, corrected, and exported as .avi extension files. The nonspecific fluorescence decay was corrected using Fiji and default options using the bleaching correction tool. Videos were digitally enhanced with Fiji–implemented filters, correcting noise using the Gaussian blur option and pixel width set to 1.0. smFISH images were corrected using Dust and Scratches filter with pixel width set to 1.0 and threshold to 0. Levels were corrected in the RGB channel and mid–tones were adjusted to increase the signal–to–noise ratio.

### Data availability and large–scale data sets

The RNA sequencing data of the T–RIP and total RNA datasets have been submitted to the BioStudies database (Sarkans et al., 2018) under the accession number S– BSST1096 and can be accessed through the following link: https://www.ebi.ac.uk/biostudies/studies/S-BSST1096?key=58afa48a-c6c2-4366-b059-d2504392a2b2 Python scripts may be retrieved from https://github.com/Nwntastz/RNA_proxitome_paper.

## Acknowledgments

We thank Rupert Frei, Morten Petersen, Ping He, Annie Marion–Poll, Shu–Hsing Wu, and Jan Petrášek, for kindly providing published materials and Fanourios Mountourakis from Moschou’s lab for fruitful discussions. This work was supported, by the Carl Tryggers Foundation (P.N.M.), and partly by a Vetenskapsrådet (VR) research council (P.N.M), FORMAS research council (P.N.M.), the EU Marie Curie–RISE (“PANTHEON,” project number 872969; PNM), Helge Axelsson Foundation (C.L.), Hellenic Foundation of Research and Innovation Ph.D. scholarships (06526; A.M. and 05947 I.H.H.), IMBB– FORTH start–up funding (P.N.M), Ministerio de Ciencia e Innovacion from Spanish Government, Grant MINOTAUR BIO2017–84066–R (FJRC and ABRL), Grant PID2020–119737GA–I00 (MCIN/AEI/10.13039/501100011033; E.G.B.), Junta de Andalucia (ProyExcel_00587; E.G.B), Vetenskapsrådet (VR) research council (2019– 05634; P.M.), and Kempestiftelserna (JCK–2011; P.M.).

## Author contributions

Conceptualization: PNM; Methodology: CL, AM, IHH, ET, VS, ABRL, BB; Investigation: CL, AM, FJRC, VS, ABRL, EGB, XM, VM, AK; Visualization: CL, AM, FJRC, PNM; Funding acquisition: CL, FJRC, EGB, PNM; Project administration: PNM, PM, PFS, EGB; Supervision: PNM; Writing – original draft: PNM; Writing – review & editing: CL, AM, RG, EGB, VM, PFS.

## Supplemental Data files

Supplemental Table 1. Primers lists.
Supplemental Figures 1–16 (legends below figures) Supplemental Files 1-10.
  File 1. RNA-binding proteins enriched in T-RIP under NS or HS. File 2. RNAs enriched/depleted from T–RIP.
  File 3. RNA and GO terms enriched in T–RIP in both NS and HS (519 genes). File 4. RT-q-PCR RNAs used for correlation with T-RIP.
  File 5. GO terms enriched/depleted false discovery rate values. File 6. SmFISH probes used (EBF2, DFL1, RAP2.4, and PP2A). File 7. IDR levels in the APEAL dataset, both in NS and HS.
  File 8. Enrichment of *ECTs* and corresponding proteins in both APEAL and T–RIP datasets.
  File 9. Polyadenylation enrichment in PBs. File 10. Enriched RNA targets of GRP7.
  File 11. Enrichments for polysome profiling RNAs.
Supplemental Statistics File
Uncropped Western Supplemental Figure 1A

